# Spliceosomal Prp8 intein at the crossroads of protein and RNA splicing

**DOI:** 10.1101/502781

**Authors:** Cathleen M. Green, Zhong Li, Olga Novikova, Valjean R. Bacot-Davis, Fenghan Gao, Nilesh K. Banavali, Hongmin Li, Marlene Belfort

## Abstract

The spliceosome is a large ribonucleoprotein complex that removes introns from pre-mRNAs. At its functional core lies the essential Prp8 protein. Across diverse eukaryotes, this protein cofactor of RNA catalysis harbors a self-splicing element, called an intein. Inteins in Prp8 are extremely pervasive and are found at seven different sites in various species. Here, we focus on the Prp8 intein from *Cryptococcus neoformans*, a human fungal pathogen. We solved the crystal structure of this intein, revealing structural homology among self-splicing sequences in eukaryotes, including Hedgehog protein. Working with the *C. neoformans* Prp8 intein in a reporter assay, we find that the biologically relevant divalent metals copper and zinc inhibit intein splicing, albeit by two different mechanisms. Copper likely stimulates reversible modifications on a catalytically important cysteine, whereas zinc binds via the same critical cysteine with a K_d_ of ~1 nM. An intein-containing Prp8 precursor model is presented, suggesting that metal-induced protein splicing inhibition would disturb function of both Prp8 and the spliceosome. These results indicate that Prp8 protein splicing can be modulated, and that this could alter spliceosome function and RNA splicing under specific conditions.

## INTRODUCTION

The spliceosome is a massive ribonucleoprotein complex that performs intron splicing, an important process for maintaining genome diversity in eukaryotes. At the heart of the spliceosome is **p**re-m**R**NA **p**rocessing factor **8** (Prp8), a large (~250 kDa) and highly conserved protein (1). Prp8 helps generate mature mRNA by coordinating critical rearrangements at the catalytic core of the spliceosome. This essential protein has been implicated in human disease (2, 3), is evolutionarily linked to group II introns (4, 5), and is structurally related to telomerase (6). Recent advances in structural biology have shed new light onto both Prp8 and the spliceosomal machinery at atomic resolution, unveiling an unprecedented level of detail into the molecular steps of intron splicing (5, 7–12).

A particular reason for our interest in Prp8 is that across several organisms, this large protein contains a self-splicing intein at different positions, implying independent acquisition. Inteins are *in*ternal pro*teins* that invade at the DNA level and undergo transcription and translation with the host gene (13–15). The intein-containing precursor undergoes protein splicing, a process that excises the intein and ligates the flanking sequences, called exteins, to form the functional protein. Inteins are often bipartite, encoding a splicing domain for excision and ligation, and an endonuclease domain for homing (16, 17). Since some inteins are mobile, they are generally considered selfish genetic elements, but new research indicates that inteins can post-translationally regulate proteins (18–25).

Inteins are found in all three domains of life, and are especially abundant in bacteria and archaea (26). In eukaryotes, inteins are sparse, but have been found in nuclear and chloroplast genomes with distinct patterns of insertion (27). Nuclear inteins tend to be in proteins that are involved in energy metabolism and RNA processing, whereas chloroplast inteins are found in proteins that carry out transcription and replication. Out of all the intein-harboring proteins in eukaryotes, Prp8 is overwhelmingly favored. There are over one hundred inteins identified across various sites of Prp8 in different species.

Pathogenic fungi seem to be enriched for inteins (27, 28). Several notable human pathogens contain Prp8 inteins, including *Aspergillus fumigatus, Histoplasma capsulatum*, and *Cryptococcus neoformans*. Intriguingly, many organisms with Prp8 inteins also tend to be intron-rich (29). The presence of inteins in Prp8 and the correlation with intron density beg the question of an intein benefit to the host, and especially to pathogens. To begin to answer this question, we focus on the Prp8 intein from *C. neoformans*. This is a mini-intein, naturally lacking the homing endonuclease domain, at only 171 amino acid residues. The intein is also found at a highly conserved site at the center of Prp8, and thus is at the core of the spliceosome (1, 5, 30).

Studying the Prp8 intein present in *C. neoformans* addresses questions of conditional protein splicing in an important human pathogen in an entirely new domain of life. Solving the Prp8 intein structure set the stage for beginning such studies and provided evolutionary context, by revealing similarities to the metazoan Hedgehog protein. Biochemical experiments then showed that the *C. neoformans* Prp8 intein is differentially responsive to copper and zinc, metals encountered by pathogens in immune cells during infection. Both metals inhibit protein splicing, largely through a critical, catalytically active cysteine. These results suggest that inteins in eukaryotes might also be post-translational sensors, echoing an emerging theme of cysteine-based regulation observed in bacterial and archaeal systems. Further, creation of a Prp8 precursor model illustrates how intein presence relates to the host protein and hints at how the intein could influence spliceosome dynamics and RNA splicing.

## RESULTS

### Prp8 is an intein hot-spot with diverse insertion sites

Recent data mining revealed over 100 inteins in the Prp8 protein. These Prp8 inteins are present across assorted eukaryotic groups, some of which emerged as far back as ~1,100 million years ago (Mya) (Fig. 1A, left) (27, 31–33). The vast majority of Prp8 inteins are found across different fungal species, particularly in Ascomycota, and the rest are dispersed across other eukaryotic phyla (Fig. 1A, left). To characterize these diverse Prp8 inteins, we performed comparative and phylogenetic analyses on a representative subset based on the splicing motifs (Fig. 1; Figs. S1 and S2) (15, 34). In total, there are seven distinct intein insertion points, denoted Prp8-**a** through Prp8-**g** (Fig. 1; Fig. S1) (33, 35, 36). With only a few exceptions, fungal Prp8 inteins occupy the same insertion site, Prp8-**a** (Figs. 1A and 1B) (27, 31, 33). The newest insertion site, Prp8-**g**, was found at the N-terminal end of Prp8 in the social amoeba *Acytostelium subglobosum* (37) and is reported here for the first time (Fig. S1B).

**Figure 1.**
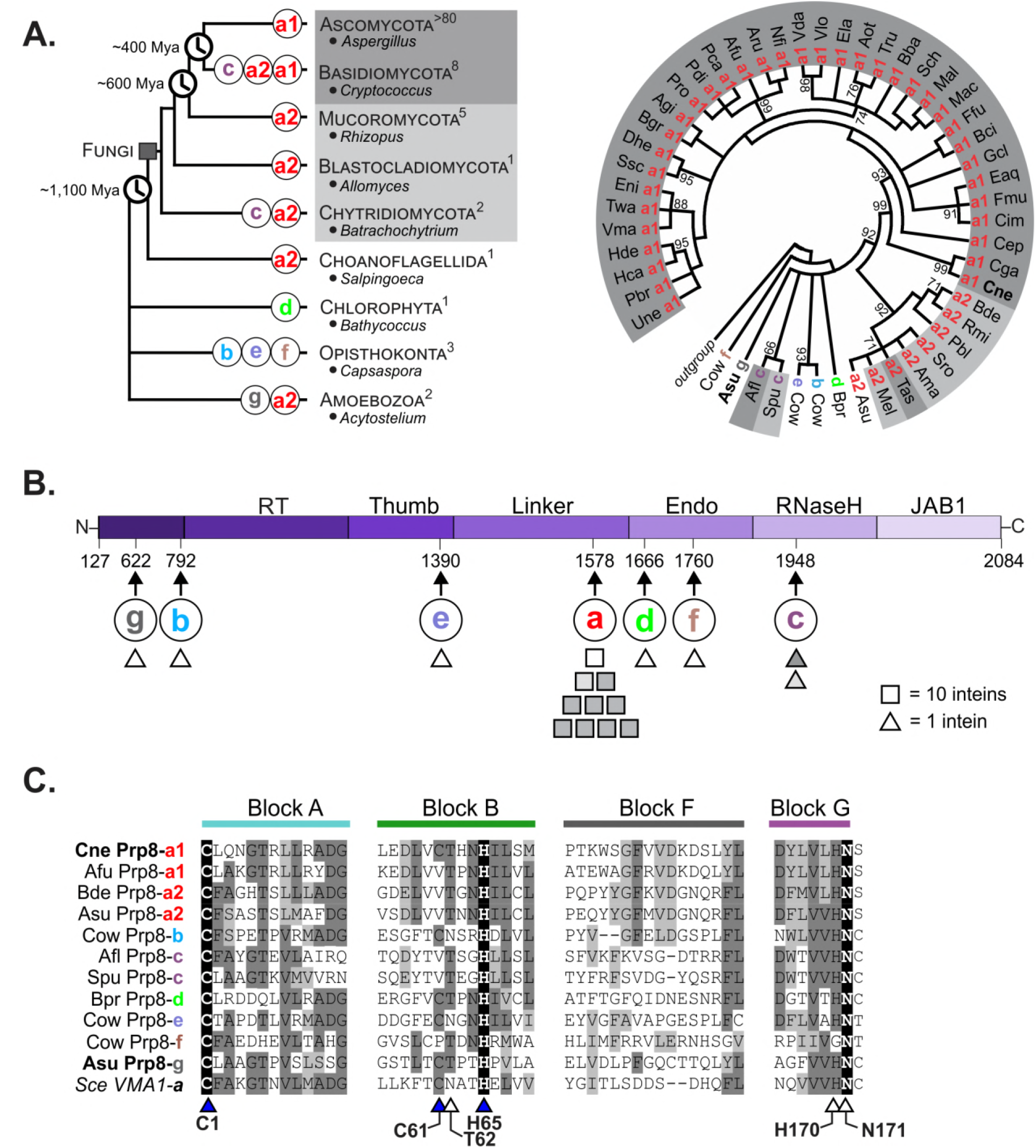
Prp8 is an intein hot-spot with multiple, independent insertion sites. **A**. Modified phylogenetic tree of eukaryotes (left) shows the phyla that contain Prp8 inteins, with representative genera listed. Evolutionary divergence times are denoted in millions of years ago (Mya). The number of Prp8 inteins in each phylum is shown in superscript, and insertion sites are shown on the branches (colored **a-g**). A phylogenetic intein tree (right) was reconstructed based on the amino acid sequences of intein splicing blocks for a subset of 50 Prp8 inteins. The radial tree shows numerous clusters, which correspond to grouping by insertion site. Abbreviated species names are shown (full names in Fig. S1A). Shading (dark gray, light gray, or white) indicates phylogenetic distribution. The divergence of the inteins despite extein conservation suggests independent invasions. **B**. A line diagram of Prp8 exteins shows the domains of the host protein from amino acid residues 127 to 2084. The arrows below indicate the site of intein insertion (**a-g**) with the corresponding residue number based on *Saccharomyces cerevisiae* Prp8 (PDB 5GMK, chain A). Shapes represent how many inteins are found at each site (square = 10 inteins, triangle = 1 intein) and are shaded to denote phylogenetic origin as in Fig. 1A. Prp8-**a** is the most common insertion site with ~100 inteins. **C**. Multiple sequence alignment of the splicing blocks of Prp8 inteins from each insertion site. Comparative analysis of residues found in blocks A, B, F, and G reveals that Prp8 inteins occupying other insertions sites are substantially different from one another, indicating independent acquisition. Identical residues are critical to self-splicing. Triangles indicate residues of general interest and those shaded blue are of specific interest. Numbers correspond to the *Cryptococcus neoformans* (*Cne*) Prp8 intein. Shading is as follows: black – identical amino acid, dark gray – conserved amino acid, light gray – similar amino acid substitution.

The reconstructed phylogenetic tree reveals that Prp8 inteins group by insertion site (Fig. 1A, right; Fig. S1A). Although insertion sites **b** through **g** have limited representation, the observed clustering, as well as the level of sequence divergence between inteins from different insertion sites, suggests multiple independent intein invasion events throughout evolutionary history. Importantly, bifurcation of Prp8-**a** inteins into two, well-supported clusters (**a1** and **a2**, with a bootstrap value of 92%) indicates recurrent invasion of inteins into site **a** across diverse fungi. All seven insertion sites were mapped to a simplified line diagram of Prp8 and are peppered across the various domains (Fig. 1B).

A multiple sequence alignment of the intein splicing motifs, referred to as blocks A, B, F, and G, demonstrates the sequence divergence among Prp8 inteins (Fig. 1C; Fig. S2). Other than identical residues located in blocks A, B, and G (Fig. 1C, black shading; Fig. S2), Prp8 inteins share limited sequence homology. Block A contains the first residue of the intein, a highly conserved cysteine called C1, which performs the first nucleophilic attack of the protein splicing pathway. This amino acid is identical across the disparate Prp8 inteins (Fig. 1C, Block A; Fig. S2). The B block usually carries a highly conserved motif known as TxxH (38). Across the Prp8 inteins, the B block histidine of TxxH is present in all analyzed inteins, whereas the threonine is somewhat conserved (Fig. 1C, Block B; Fig. S2). Also alike across all Prp8 inteins is a terminal asparagine at the C-terminus of the intein in block G, which also participates in splicing (Fig. 1C, Block G; Fig. S2). The first amino acid of the C-extein, known as the +1 residue, is usually a cysteine, serine, or threonine, and all Prp8 inteins use one of these as the +1 nucleophile. The F block shows little conservation across Prp8 inteins. The *Saccharomyces cerevisiae* (*Sce*) VMA1 intein, in the vacuolar ATPase, is as similar to Prp8 inteins as other Prp8 inteins are to each other (Fig. 1C), indicating a close ancestral relationship. The poor sequence alignment among Prp8 inteins reinforces that distinct inteins recurrently invaded Prp8.

### *C. neoformans* Prp8 intein structure shows homology to eukaryotic self-splicing elements

We next solved the crystal structure of the *Cryptococcus neoformans* (*Cne*) Prp8 intein found at site **a** (Fig. 2A). This intein was chosen due to its small size, and because it is found in a significant human pathogen. The *Cne* Prp8 intein was solved to 1.75 Å resolution (Fig. 2A). This novel structure helped us to develop a sense of structural relatedness of the *Cne* Prp8 intein to other inteins and intein-like elements, and to later model the intein into both Prp8 and the spliceosome.

**Figure 2.**
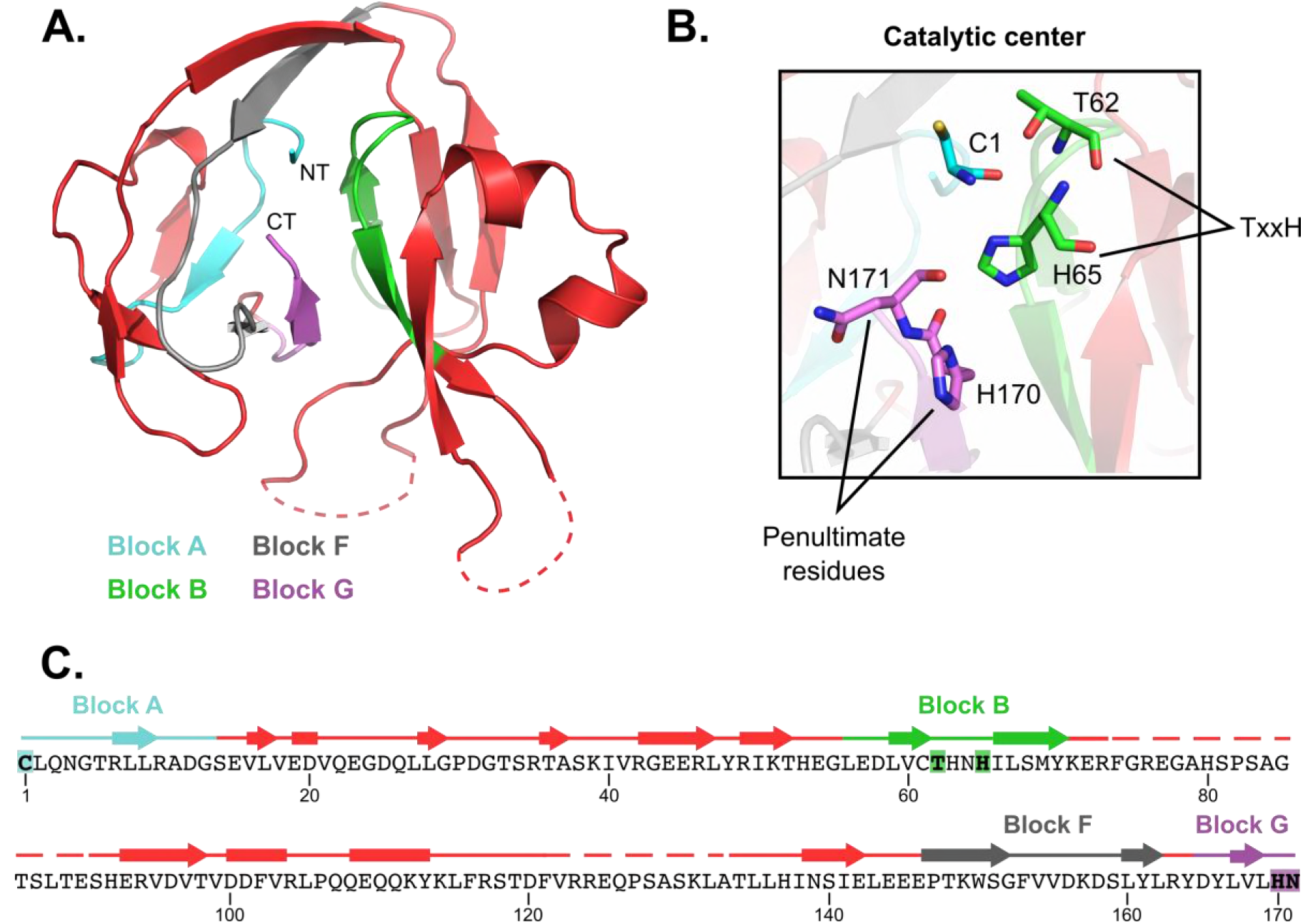
Structure and features of the *Cne* Prp8 intein. **A.** A crystal structure of the *Cne* Prp8 intein from site **a** was solved to 1.75 Å resolution. This structure has the canonical horseshoe shape and resolved the four splicing blocks, indicated in cyan (block A), green (block B), gray (block F), and purple (block G) (Fig. 1C). The amino (NT) and carboxy termini (CT) are annotated. Some regions of the structure are unresolved (dashed lines), and likely represent remnants of the original endonuclease domain or linker sequences. **B.** Features of the active site. The catalytic center is shown, highlighting the first residue (C1) and the penultimate residues (H170 and terminal N171). The B block TxxH motif is shown in green with the T62 and H65 shown as sticks. These residues are critical to carrying out autocatalytic protein splicing. **C.** Sequence of the *Cne* Prp8 intein (residues 1-171) overlaid with secondary structure features. Blocks distant in sequence fold close in 3D space to allow protein splicing to occur. Unresolved regions (dashed lines) are between blocks B and F. Arrows represent β-strands, rectangles are α-helices. Residues noted in Fig. 2B are highlighted.

The *Cne* Prp8 intein structure represents only the second known fungal nuclear intein structure. The first was of the *Sce* VMA1 intein, which was solved with its linker domain, a connector between the splicing blocks and the internal endonuclease domain (39, 40). As with all solved intein structures so far, the *Cne* Prp8 intein has the canonical horseshoe shape, created by pseudo-two-fold symmetry that positions the catalytic N- and C-termini in close proximity (Fig. 2A). Highlighting the splicing blocks (A, B, F, and G), we see the active core that carries out autocatalytic protein splicing (Figs. 2A and 2B) (15, 34). This catalytic center contains the residues essential for splicing: the nucleophilic cysteine, called C1, in block A, and the terminal asparagine (N171) in block G (Fig. 2B). The C1 and N171 are also positioned in the vicinity of the conserved B block TxxH residues (T62 and H65), which are important for priming the intein for autocatalytic excision at its amino terminus (Fig. 2B) (41). All of these residues contribute critically to the protein splicing pathway, which involves a series of nucleophilic attacks, cyclization of the terminal asparagine, and reformation of a peptide bond between the exteins to form the functional protein (42). Overlaying the *Cne* Prp8 intein primary sequence with its secondary structure shows the position of residues from each block within the context of the 3D architecture (Fig. 2C). For example, blocks A and B are far apart in sequence, but fold proximally in 3D space to execute protein splicing (Figs. 2B and 2C). This representation also illustrates that the unresolved regions of the intein are between block B and block F, and likely represent flexible linker sequences of a former endonuclease domain (Fig. 2C).

We next performed a 3D BLAST to compare the *Cne* Prp8 intein structure to other solved structures. Unsurprisingly, the *Cne* Prp8 intein structure as the query returns the *Sce* VMA1 intein as the top hit (Fig. 3A, PDB 1GPP) (39). These are both fungal inteins encoded in nuclear genomes. An overlay of the *Cne* Prp8 intein (red) and the *Sce* VMA1 intein (splicing domain in cyan, linker/endonuclease domain in gray) displays remarkable structural similarity in the splicing modules (Fig. 3A, RMSD of 1.06 Å). The unstructured regions in the *Cne* Prp8 intein structure are where the *Sce* VMA1 intein encodes a linker domain (Fig. 3A, dashed red lines). A closer look at the active centers of the *Cne* Prp8 intein and the *Sce* VMA1 intein demonstrates unmistakable overlap of the catalytic residues (Fig. S3A), further confirming the similarities.

**Figure 3.**
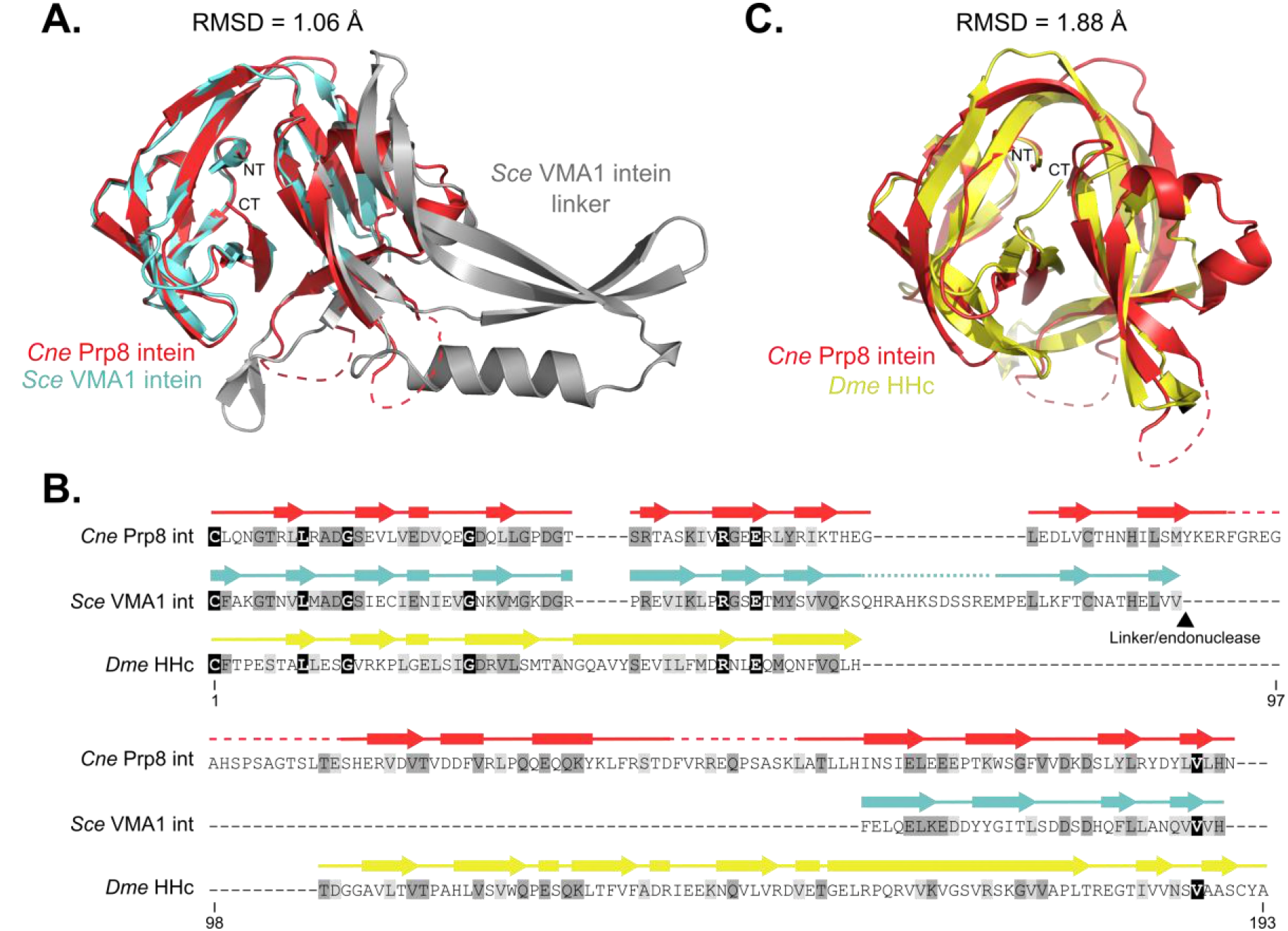
Homology of the *Cne* Prp8 intein to eukaryotic self-splicing elements. **A.** Overlay of the *Cne* Prp8 intein (red) with the *Sce* VMA1 intein (cyan and gray, PDB 1GPP) with an RMSD of 1.06 Å. Structural similarities are most pronounced in the splicing domain (blocks A, B, F, and G) of the *Sce* VMA1 intein, whereas the linker/endonuclease domain (gray) is where the *Cne* Prp8 intein did not resolve (dashed lines). The amino (NT) and carboxy termini (CT) are annotated. **B.** Multiple sequence alignment of *Cne* Prp8 intein, *Sce* VMA1 intein, and *Drosophila melanogaster* (*Dme*) HHc. Sequence comparison (residues 1-193) reveals significant differences across the three proteins, which are only 22.6% identical. Overlaying secondary structure shows that, despite sequence divergence, the proteins have high structural similarity. Shading is as follows: black – identical amino acid, dark gray – conserved amino acid, light gray – similar amino acid substitution. Arrows represent β-strands, rectangles are α-helices. Unresolved regions shown as dashed lines. **C.** Overlay of the *Cne* Prp8 intein (red) with *Dme* HHc (yellow, PDB 1AT0). Structural 3D BLAST shows parallels between the eukaryotic intein and the eukaryotic self-splicing Hedgehog domain, with an RMSD of 1.88 Å. The amino (NT) and carboxy termini (CT) are annotated.

Remarkably, another top hit from the 3D BLAST is the 17 kDa fragment of the *Drosophila melanogaster* (*Dme*) Hedgehog C-terminal domain (HHc) (43). Hedgehog is an essential signaling molecule in higher eukaryotes with an analogous self-cleaving reaction performed by a highly conserved cysteine (44). The cleavage allows the N-terminal domain of Hedgehog to ligate to a cholesterol molecule, which plays a critical role in metazoan development. There has been considerable speculation about the relatedness of Hedgehog and inteins (43, 45). It was recently reported, based on sequence similarity, that the N-terminal portion of Hedgehog was acquired through horizontal gene transfer from a prokaryote (46). However, a sequence alignment between the *Cne* Prp8 intein, the *Sce* VMA1 intein, and *Dme* HHc shows the three eukaryotic self-splicing sequences have only an average of 22.6% sequence identity (Fig. 3B, 26.4% identity VMA1 to Prp8, 19.2% HHc to Prp8, 22.2% VMA1 to HHc). One highly conserved residue across the three proteins is the initiating cysteine, shared by all the sequences, as well as a C-terminal valine (Fig. 3B, black shading). Despite the considerable sequence divergence, a secondary structure overlay demonstrates that these sequences all code for the same structural elements (Fig. 3B). Strikingly, the *Cne* Prp8 intein and *Dme* HHc have an RMSD of 1.88 Å (Fig. 3C, PDB 1AT0), sharing a similar degree of structural relatedness as a bacterial and a fungal intein (Fig. S3B, RMSD 2.22 Å, PDB 2IMZ) (47). These results reinforce the evolutionary connection between inteins and Hedgehog proteins.

### *C. neoformans* Prp8 intein is responsive to stress

With the structure solved, we next sought to investigate Prp8 intein splicing and if it is regulated in any way. For simplicity, the *Cne* Prp8 intein was studied in *Escherichia coli*. Given that full-length Prp8 contains ~2500 amino acids, we cloned the *Cne* Prp8 intein into a reporter construct that uses maltose binding protein (MBP) and green fluorescent protein (GFP) as foreign N- and C-exteins, respectively (21, 24) (Fig. 4A). From this construct, termed MIG, which contains five native N- and C-extein residues, expression is induced and splicing products, such as ligated exteins (LE), are visualized using non-denaturing SDS-PAGE and scanning for GFP fluorescence (Fig. 4A, left). Off-pathway C-terminal cleavage (CTC) is also detectable in the gels, and is caused when the terminal asparagine (N171) cyclizes prior to the first step of protein splicing (Fig. 4A, right).

**Figure 4.**
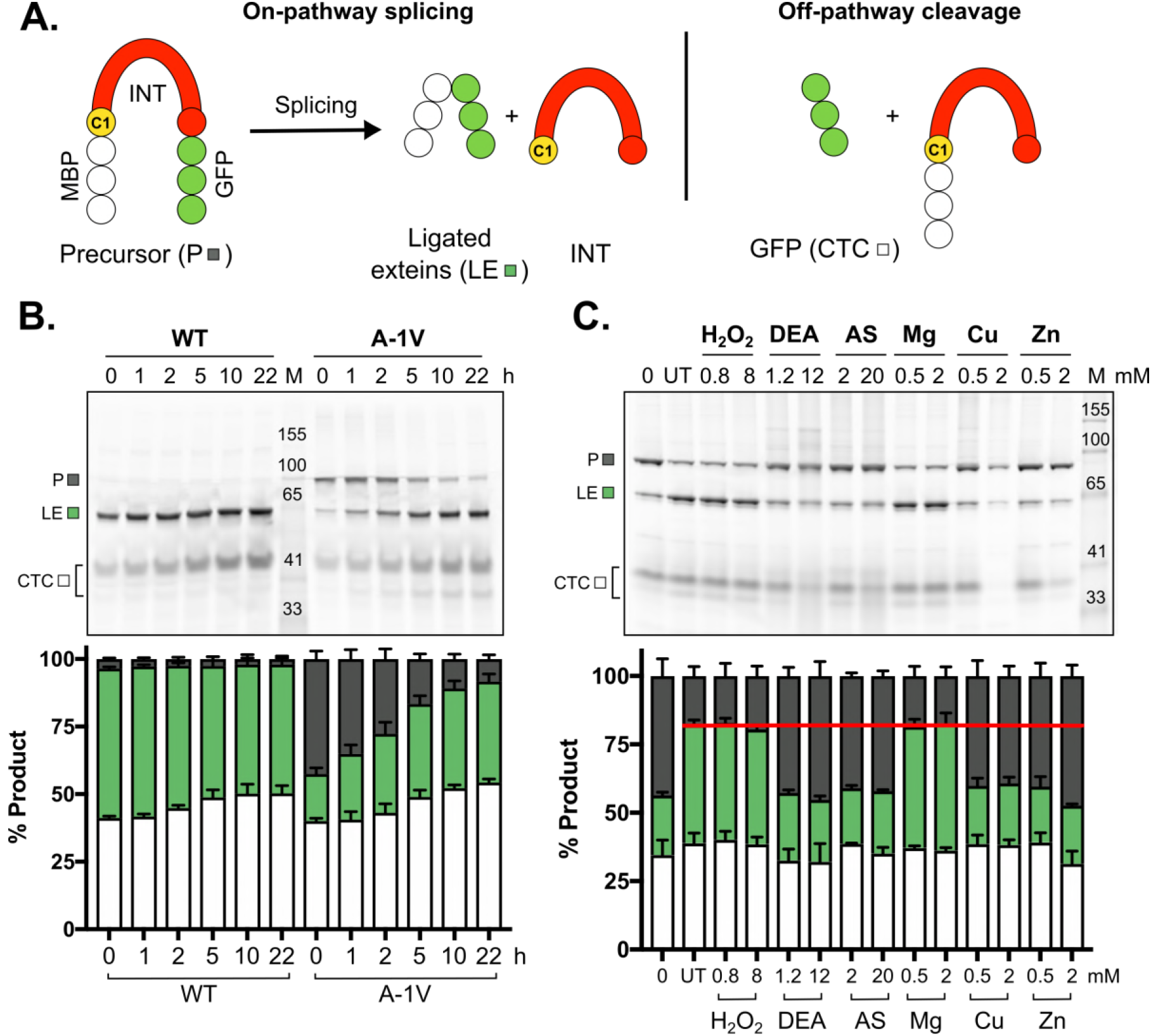
MIG Prp8 A-1V is responsive to metal and RNS treatment. **A.** Schematic of the MIG reporter. The construct contains <u>M</u>BP-Intein-<u>G</u>FP (MIG) and is expressed in *Escherichia coli*. GFP allows monitoring of splicing using in-gel fluorescence. The precursor (P) can undergo protein splicing (left), yielding ligated exteins (LE) and free intein (INT, not seen on gels), or may undergo off-pathway cleavage reactions (right), such as C-terminal cleavage (CTC). The catalytic cysteine, C1, is labeled. **B.** MIG Prp8 WT splices rapdily. A fluorescent gel of a splicing time-course shows that the wild-type *Cne* Prp8 intein in MIG is entirely spliced by the start of the assay (left, WT, 0 h). The A-1V mutant had precursor at the assay start and spliced over time (right, A-1V, 0 h). Quantitation is shown below in stacked plots. **C.** MIG Prp8 A-1V accumulates precursor under RNS and metal treatment. After 5 h *in vitro* treatment with a panel of environmental stressors, there was an increase in P compared to the untreated (UT, red line) with the RNS compounds DEA NONOate (DEA) and Angeli’s salt (AS), and the metals copper (Cu), and zinc (Zn). H_2_O_2_ and magnesium (Mg) show no effect. Quantitation is shown below in a stacked plot. Data are representative of three biological replicates and mean standard deviations are shown. Representative gels are shown.

First, we observed that the *Cne* Prp8 intein splices well in the foreign context to yield ligated exteins. However, splicing was so rapid that the amount of precursor remaining after induction (0 h) did not provide a suitable dynamic range for performing splicing assays (Fig. 4B, WT). To slow down splicing and accumulate precursor, mutations were made to the last residue of the N-extein (referred to as the −1), a site previously shown to affect splicing rates (48). After random mutagenesis, a slower splicing mutant was isolated (Fig. 4B, A-1V). The MIG Prp8 A-1V mutant has 40% precursor at 0 h and is splicing active over time (Fig. 4B, A-1V). It is worth noting that splicing rates are intein-dependent, given that other Prp8-**a** inteins from fungal pathogens exhibit diverse splicing phenotypes when cloned into MIG (Fig. S4).

Next, using MIG Prp8 A-1V, we asked if a condition exists in which intein splicing might be regulated. Treatments chosen were to mimic environmental stress that *C. neoformans* experiences during infection, such as reactive oxygen species (ROS), reactive nitrogen species (RNS), and metals, all of which prevail during the intracellular respiratory burst (Fig. 4C) (49, 50). From this initial panel, the RNS compounds DEA NONOate and Angeli’s salt showed significant precursor accumulation (Fig. 4C, DEA and AS). It also appears that copper and zinc can cause splicing inhibition of MIG Prp8 A-1V (Fig. 4C, Cu and Zn). Under these conditions, splicing was inhibited by ~50% (Fig. 4C). This preliminary compound screen indicates that the *Cne* Prp8 intein may be subject to inhibition by specific conditions that occur during infection.

### Splicing inhibition is mechanistically distinct under copper and zinc treatment

Metal binding has been reported for other inteins, and often engages catalytic residues, which would stall protein splicing (47, 51–56). Therefore, we chose to follow up on the observed copper and zinc inhibition by running *in vitro* MIG Prp8 A-1V assays to further assess effects on protein splicing over time (Fig. 5).

**Figure 5.**
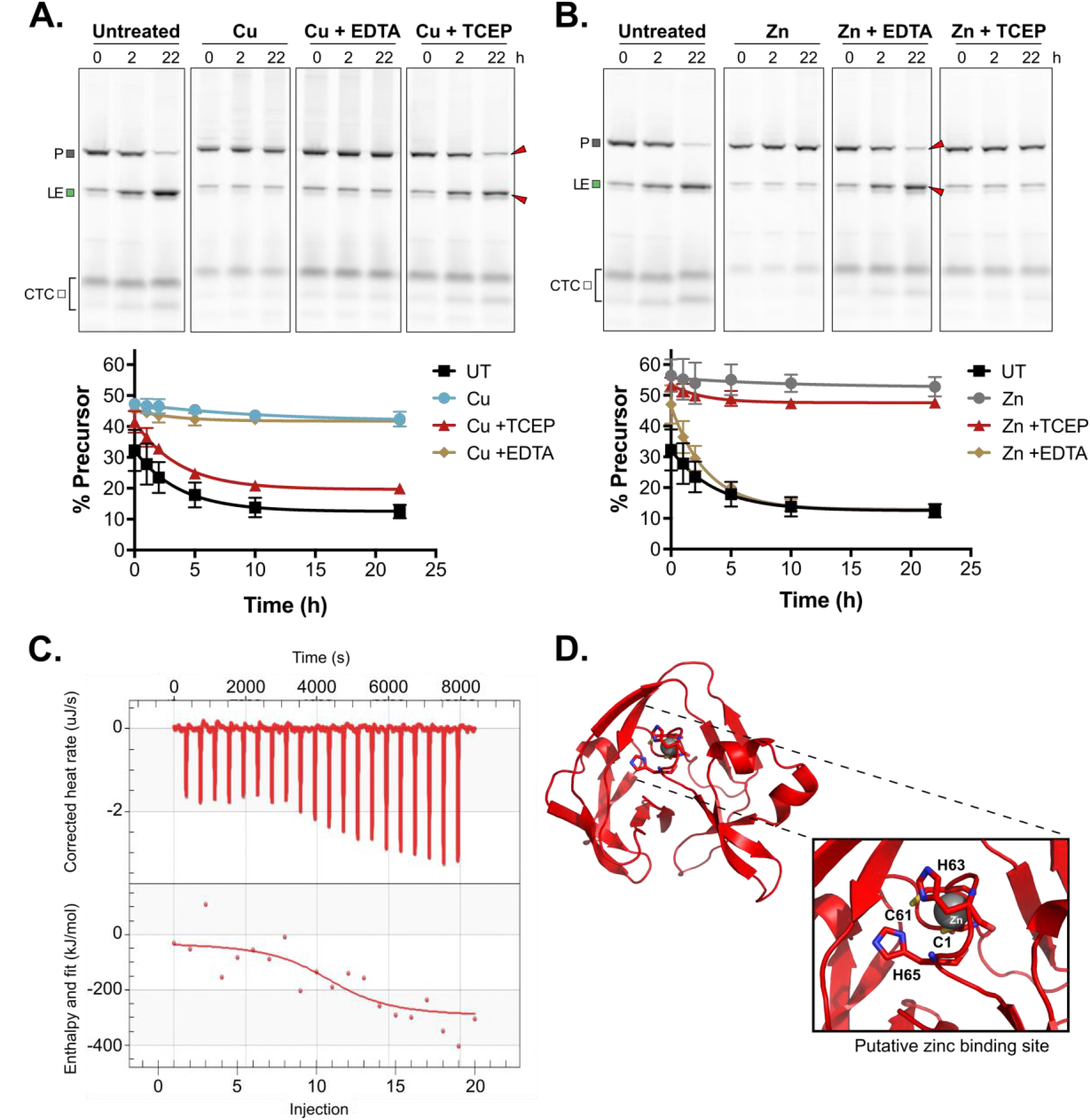
MIG Prp8 A-1V is differentially inhibited by copper and zinc. **A.** Copper inhibition is reversible by reducing agent only. MIG Prp8 A-1V splicing was completely inhibited by copper treatment (Cu) over 22 h, as minimal loss in P or increase in LE occurred compared to the untreated control. The inhibition was unaffected by treatment with metal chelator EDTA (Cu + EDTA). Upon adding copper and then reducing agent TCEP (Cu + TCEP), splicing was restored and P converted into LE over time. Red arrows indicate splicing rescue. The splice products were quantitated and the percent precursor is plotted as a proxy for splicing inhibition. Representative gels are shown. **B.** Zinc treatment is reversible by EDTA only. MIG Prp8 A-1V splicing was strongly inhibited by zinc treatment (+ Zn) over 22 h compared to untreated lysates. The zinc-based inhibition reversed when treated with EDTA (Zn + EDTA) and splicing was observed at a rate comparable to the untreated samples. Red arrows indicate splicing rescue. When adding zinc and then reducing agent TCEP (Zn + TCEP), splicing was unaffected. Plots are as in A above. Data are representative of three biological replicates and mean standard deviations are shown. Trend lines are fit to show the decay curve. Representative gels are shown. **C.** Zinc binds to the *Cne* Prp8 intein tightly. Using isothermal titration calorimetry (ITC), 16 μΜ purified *Cne* Prp8 intein was titrated with 0.05 mM ZnSO_4_ over 20 injections at 37°C and pH 7.0 on a Nano TTC. The binding isotherm (bottom) shows integrated heat per mole of ZnSO_4_ as a function of the molar ratio of ZnSO_4_ to the *Cne* Prp8 intein and a K_d_ ~1 nM was calculated (Table 1). The NanoAnalyze TTC software automatically discarded outlier data points. Experiment was performed in triplicate. **D.** A putative zinc-binding pocket in the *Cne* Prp8 intein. Zinc binding was modeled using a structure of zinc bound human ubiquitin ligase (PDB 5TDA) (87) and the *Cne* Prp8 intein structure. A coordination site is proposed using C1, C61, H63 and H65 (shown as sticks), with one zinc ion binding, shown as a gray sphere.

We found that 1 mM CuSO_4_ caused strong splicing inhibition for up to 22 h compared to untreated controls (Fig. 5A, Untreated and Cu). This inhibition persisted up to 30 h (Fig. S5A). To test a copper-binding hypothesis, the same assay was carried out, but after 2 h of incubation with copper, ethylenediaminetetraacetic acid (EDTA) was added in excess. EDTA chelates copper and should strip bound copper from the *Cne* Prp8 intein so that splicing can occur. However, addition of EDTA did not rescue splicing, ruling out the possibility of inhibition purely by copper binding (Fig. 5A, Cu + EDTA).

Copper is a redox active metal that can cause cysteine oxidation, either by promoting disulfide bond formation or by catalyzing reversible or irreversible oxidative modifications (57). We next tested whether the *Cne* Prp8 intein cysteines are being reversibly modified by copper, which would prevent the C1 from performing the first nucleophilic attack, and has precedent in intein biology (21, 25). We added the reducing agent tris-(2-carboxyethyl)phosphine (TCEP) to the lysates after a 2 h incubation with copper. Strikingly, TCEP completely reversed the splicing inhibition (Fig. 5A, Cu + TCEP). After reduction, MIG Prp8 A-1V precursor conversion into ligated exteins occurred at a rate similar to that of no copper treatment (Fig. 5A, bottom), indicating reversible cysteine oxidation.

The *Cne* Prp8 intein only has two cysteines: C1, and C61 in the B block, immediately preceding the TxxH motif (see Fig. 1C, blue arrowheads). The C1 to C61 distance is 8.9 Å, too far for a disulfide bond to form (Fig. S5B), although, because of residue flexibility, this does not strictly preclude formation of a disulfide (21, 58, 59). We also found that C61 is not highly conserved across Prp8 inteins (Fig. S5C), and the most commonly used residue at this site is valine. Therefore, to ask if C1 modifications are sufficient to inhibit protein splicing, several mutants of C61 in MIG Prp8 A-1V were tested for splicing activity (Fig. S6A) and treated with copper (Fig. S6B). The C61 mutants also showed precursor accumulation (Fig. S6B), suggesting that C1-C61 disulfide bonding is not the underlying inhibitory mechanism, and that copper induces at least C1 oxidation, which is enough to cause the non-splicing phenotype.

We further confirmed cysteine modification by performing mass spectrometry on purified *Cne* Prp8 intein. This showed a peak shifted by 32 Da, consistent with an addition of two oxygen atoms (Fig. S7A). Additional validation pinpointed reversible sulfenic acid modifications (-SOH) to C1 and C61 with copper treatment (Fig. S7B), but these were also present in the untreated *Cne* Prp8 intein (Fig. S7A). This indicates that the *Cne* Prp8 intein has highly reactive cysteines that can be modified by atmospheric oxygen alone. Such extreme sensitivity has been observed for other inteins that are regulated by cysteine modification (21). At this time, it is unclear whether the modifications in this assay are the result of copper, oxygen in air, or both. Based on our MIG data, reversible, copper-induced cysteine modifications are the most likely explanation for the inhibition we observe (Fig. 5A), and are likely mediated mainly through C1 (Fig. S6).

Next, zinc, a metal without redox activity, was added to MIG Prp8 A-1V lysates given that it too was inhibitory in preliminary treatments (Fig. 4C). The addition of 1 mM ZnSO_4_ also caused protein splicing inhibition, and for similar time periods (Fig. 5B, Untreated and Zn). To probe the mechanism of zinc inhibition, we followed up with the same EDTA chelation and TCEP reduction. In contrast to copper, EDTA reversed protein splicing inhibition with zinc (Fig. 5B, Zn + EDTA), but reduction did not (Fig. 5B, Zn + TCEP). Thus, zinc likely causes inhibition by binding to the *Cne* Prp8 intein, as it is redox inactive and TCEP treatment yielded no change.

To corroborate zinc binding, purified *Cne* Prp8 intein was titrated with zinc in an isothermal titration calorimetry (ITC) experiment. This revealed tight binding of zinc to the intein, with a Kd in the 1 nM range, whereas no copper binding was detected in a biologically relevant range (Fig. 5C, Table 1). To further understand the mechanism of zinc binding, we turned to molecular modeling using the crystal structure to predict a coordination site. The model was built by surveying published zinc-bound structures, assessing the composition of the zinc pockets, and looking for similarities in the *Cne* Prp8 intein. The putative binding site uses two cysteines and two hisitidines, a common tetrahedral geometry for a non-catalytic zinc coordination (60). Specifically, we propose that C1, C61, and the two B block histidines, H63 and H65, shift slightly to coordinate a zinc ion (Fig. 5D; Figs. S8A and S8B). These residues are all in or nearby critical splice sites, which would explain the splicing inhibition. To test if this is indeed the zinc site, we performed ITC with individual mutants C1A, C61A, H63A, and H65A and a triple C1A/C61A/H65A mutant. This demonstrated a significant decrease in binding, with K_d_ values increased by 500-fold or more, supporting the putative zinc pocket in the *Cne* Prp8 intein (Fig. S8C, Table 1). We confirmed the mutants are structurally sound using 1D proton NMR, so the observed decrease in binding is likely due to loss of a coordination residue (Fig. S8D).

**Table 1.**
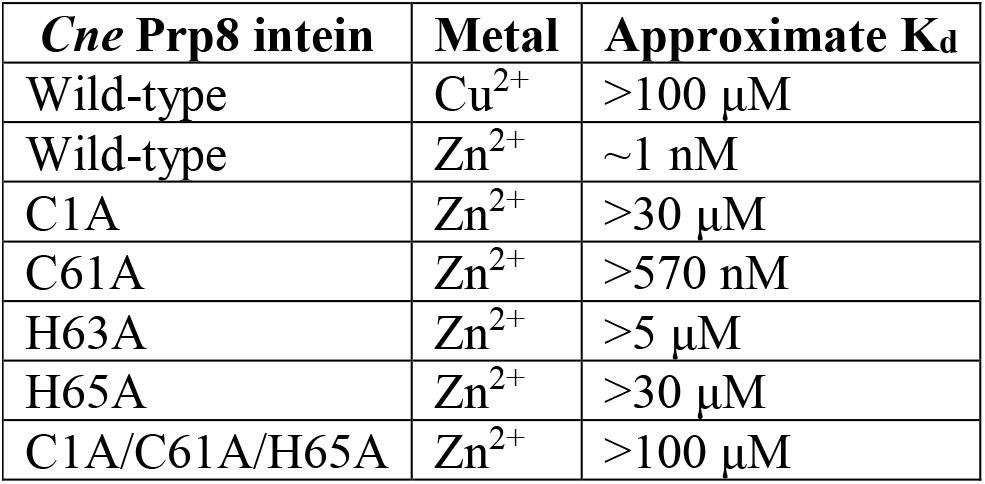
Metal binding K_d_ estimates using ITC

### A precursor model of Prp8 relates intein retention to spliceosome function

Finally, we wished to determine how protein splicing inhibition might affect both Prp8 and the spliceosome. Therefore, we docked the intein into a known Prp8 structure and generated a precursor model, where the intein is still covalently connected to the exteins. In this model, the bonds flanking the intein were broken at site **a** in Prp8 from a cryo-EM C complex spliceosome solved from *Schizosaccharomyces pombe* (Fig. 6A, PDB 3JB9, chain A) (12). The *Cne* Prp8 intein structure was computationally inserted using an energy optimization protocol, allowing insight into how intein presence might affect Prp8 and RNA splicing in general.

**Figure 6.**
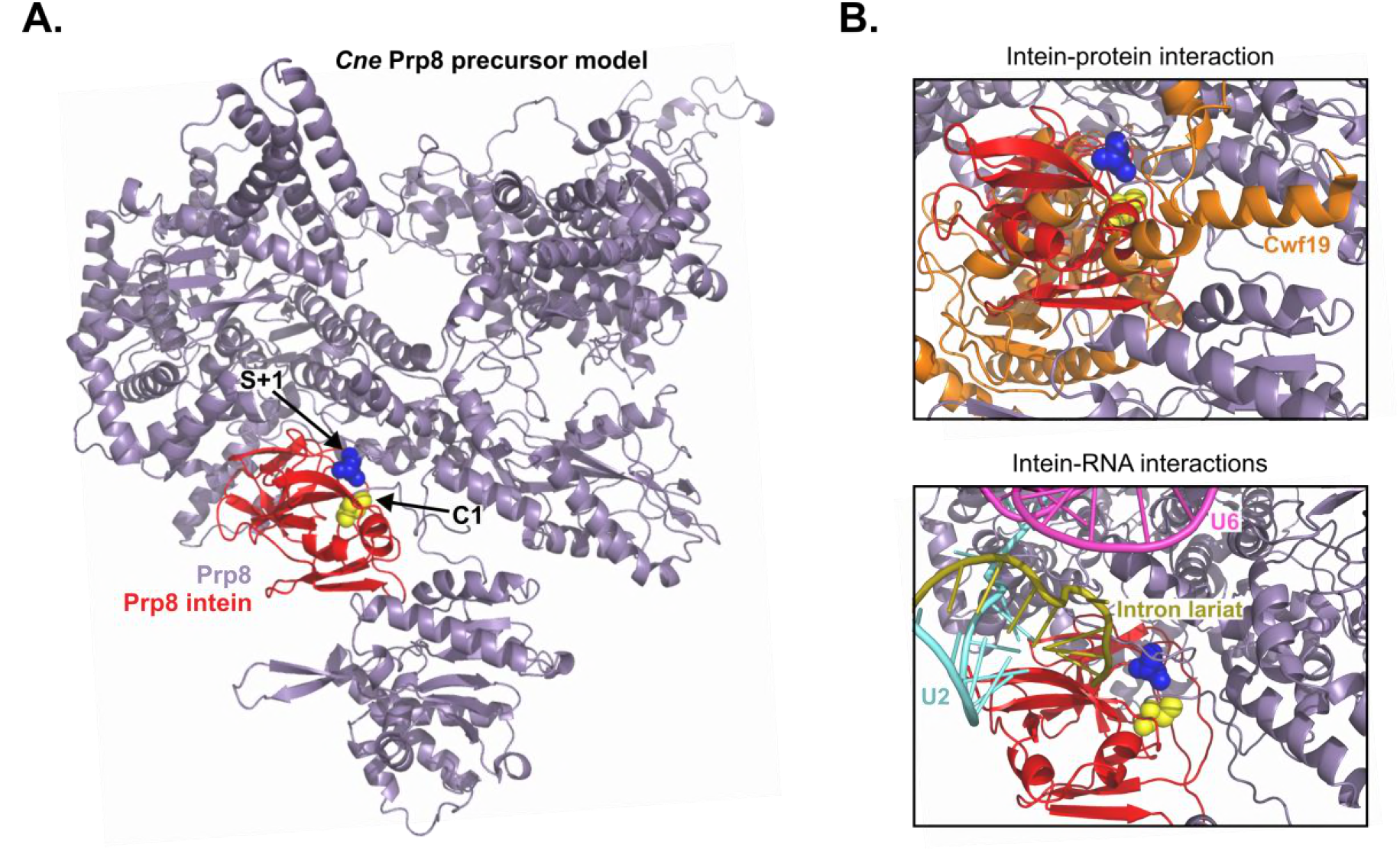
Modeling the *Cne* Prp8 intein into Prp8 and docking into the spliceosome reveals unfavorable interactions. **A.** *Cne* Prp8 intein in Prp8 exteins. The *Cne* Prp8 intein structure (red) was modeled into a structure of *Schizosaccharomyces pombe* (PDB 3JB9, chain A) Prp8 exteins (lavender). The S+1 is shown as blue spheres to indicate the site where a peptide bond was broken to insert the intein. The C1 is shown as yellow spheres and specifies start of the intein. The Prp8 intein fits into a cleft of the Prp8 structure, in a region that is highly conserved and functionally important. **B.** The *Cne* Prp8 intein in the spliceosome. The intein-containing Prp8 was overlaid into a spliceosome structure (PDB 3JB9). This revealed a direct clash of the intein with spliceosomal protein Cwf19 (top panel, orange). There are also possible clashes with the intein and the intron lariat (split pea) and U2 snRNA (teal). U6 snRNA is also in the vicinity (pink).

From our precursor modeling, it appears that the *Cne* Prp8 intein is fairly well accommodated in the Prp8 protein (Fig. 6A). Prp8 looks like a curved bean, and the intein fits comfortably into its open cleft. This insertion, site **a**, is in a linker domain that is structurally flexible, as demonstrated by poor resolution of this region in solved structures (5). Given this flexibility, the intein might be able to adopt several configurations at site **a**. The intein positioning shown here is an open orientation, and likely the least disruptive, which would allow for the Prp8 exteins to fold properly. Nonetheless, the presence of the intein likely still interrupts Prp8 function, given the importance of this region, the supporting contacts Prp8 makes within the spliceosome, and the RNA splicing defects in Prp8 mutants (1, 2) (see Discussion). Mapping the other insertion sites onto a Prp8 structure from a *S. cerevisiae* spliceosome also reveals that their presence would presumably disrupt Prp8 function, as they too cluster around the active site (Fig. S9, PDB 5GMK, chain A) (10).

We next overlaid the Prp8 intein-containing precursor in the spliceosome from *S. pombe* (Fig. 6B, PDB 3JB9) (12). Although this cleft region also occupies a relatively sparse area of the spliceosome (Fig. S10), there are a few crucial players, both protein and RNA, in the vicinity of the intein. For example, one essential splicing protein, Cwf19, occupies the same 3D space as the intein (Fig. 6B, Intein-protein). Cwf19 is implicated in displacing the intron lariat/U2 branch helix (61). Furthermore, there are a few important RNAs in the area of the intein (Fig. 6B, Intein-RNA). These include the U2 snRNA and the intron RNA close by, with U6 snRNA appearing distally. Our precursor model suggests that RNA splicing would be arrested or otherwise disturbed by the presence of the intein, particularly in the presence of environmental stressors that inhibit protein splicing (Fig. 4C).

## DISCUSSION

Here, we have shown that Prp8 inteins are widely distributed across eukaryotes and have invaded the Prp8 protein repeatedly and independently (Figs. 1A and 1B), suggestive of potential adaptation that provides an advantage to the host. The crystal structure of the *Cryptococcus neoformans* (*Cne*) Prp8 intein showed similarities to the metazoan Hedgehog protein and has facilitated studies of function, as well as provided a basis for molecular modeling (Figs. 2, 3, and 6). Initial *in vitro* studies demonstrated that some environmental stressors that are prevalent in infected macrophages are capable of modulating protein splicing of the *Cne* Prp8 intein (Fig. 4C). Specifically, copper and zinc are potent inhibitors of protein splicing, with each metal interacting with the intein in distinct ways, albeit both through the catalytic cysteine, C1 (Fig. 5). Copper likely hinders protein splicing by cysteine oxidation, and zinc inhibits by tenacious binding to the intein (Fig. 5; Figs. S7 and S8). This work supports a growing theme in intein research that underscores the reactivity of catalytic cysteines (21–23, 58, 59). We propose that the *Cne* Prp8 intein, at the nexus of protein and RNA splicing, can sense metals to pause RNA removal during stressful conditions. This is reinforced by an intein-containing Prp8 precursor model, which suggests that protein splicing inhibition would interfere with RNA splicing (Fig. 6; Fig. S10).

### Prp8 is an intein sink, with functional implications

We demonstrated a broad distribution of Prp8 inteins with multiple insertion sites (Figs. 1A and 1B), a pattern noted by others as well (27, 31, 33). Our data support the notion that Prp8 was invaded repeatedly, including at least twice at site **a** (Fig. 1A, **a1** and **a2**), and the intein retained, with functional implications. We discovered a novel insertion in a social amoeba, site **g**, bringing the total number of known insertion sites in Prp8 to seven (Fig. 1C; Fig. S1B). Similar trends were previously reported with the mycobacterial iron-sulfur cluster assembly protein SufB, which has three distinct insertion sites, and the mycobacteriophage terminase TerL, which has at least 5 intein insertion sites (20, 21, 26). Such bioinformatics observations have led to fruitful research on intein function, which is now beginning to show that inteins can be tuned to respond to environmental cues (19, 21, 23, 24). A striking example is a mycobacterial intein in DnaB helicase, located in the P-loop of the ATPase domain (62), which is sensitive to ROS both *in vitro* and *in vivo* (58).

### Structural insights into the Prp8 intein

Here, we present the second structure of a eukaryotic intein, and the sole structure of a eukaryotic intein in an essential protein (Fig. 2A). The *Cne* Prp8 intein structure provides insight into the similarity of inteins in eukaryotes (Figs. 3A-C), suggesting that they likely evolved from a common ancestor. Remarkably, the *Cne* Prp8 intein also has a comparable structure to the C-terminus of a Hedgehog protein (Fig. 3C), which executes a cleavage and ligation reaction to cholesterol also by utilizing a cysteine (44). These results suggest that eukaryotic inteins and Hedgehog proteins might be ancestrally related, but why inteins do not exist in metazoan genomes is a puzzle yet to be explained.

Around a dozen intein structures have been solved so far, comprising mainly bacterial and archaeal inteins (63). These have proven useful for studying inteins as novel drug targets (51). As inteins often invade essential proteins in pathogens, inhibiting them from splicing out is an attractive option for developing new antimicrobials (51, 64, 65). Progress towards this goal has been made in prokaryotes using the mycobacterial RecA recombinase intein. A co-crystal of the RecA intein and the antineoplastic compound, cisplatin, helped resolve the mechanism of protein splicing inhibition (51). This showed that the platinum ions of cisplatin bind to the RecA intein at its two catalytic cysteines, C1 and C+1. Concurrent work studying cisplatin and the Prp8 intein also demonstrated effective splicing inhibition, both *in vitro* and *in vivo*, although the mechanism is different than the RecA intein (Li et al., in prep). Solving the *Cne* Prp8 intein structure, along with the observed metal inhibition, is a catalyst for advancing these studies in an essential protein in a eukaryote, at an opportune time given that the antifungal pipeline is drying up (66).

### *C. neoformans* Prp8 intein is responsive to metals, with biological ramifications

Pathogenic microbes occupy niches that expose them to the opposing toxicities of metal ion excess and deprivation (67). A major reason why *C. neoformans* can successfully colonize distinct tissues is its ability to adapt to the ever-changing environments of the host (68, 69). *C. neoformans* infects the lungs and can disseminate to the brain, causing deadly cryptococcal meningitis in immunocompromised individuals (70). As an intracellular pathogen, *C. neoformans* experiences the oxidative burst of the phagolysosome, which exposes it to acute metal stress (71). Levels of copper can reach up to several hundred micromolar, leading to metal ion toxicity, while zinc concentrations are initially high, but drop with ongoing infection (67, 72, 73). These means to thwart fungal proliferation by metal depletion and compartmentalization are known as nutritional immunity (74). Pathogens prevail by devising sophisticated strategies to either acquire metals or shuttle them out of the cell (67, 71, 75). This constant tug of war between the host and pathogen forces continual evolution, and novel means of overcoming metal ion excess and depletion.

Here, we speculate that the *Cne* Prp8 intein might provide cryptococci another means to sense metals during infection. For example, a pause in protein splicing may be useful for overcoming high levels of copper or zinc (Figs. 4C and 5). Copper generates destructive ROS intermediates and can displace iron from iron-sulfur clusters (72), whereas both copper and zinc can dislodge divalent metals from other metalloprotein complexes. Like other stressors known to inhibit RNA splicing, copper and zinc would act post-translationally to block Prp8 intein splicing and inhibit spliceosome function until levels of the metals are reduced by scavenger proteins or metal transporters (71). An advantage of these post-translational strategies is reversibility and instantaneous resumption of spliceosomal function with a return to normal conditions.

Post-transcriptional programs that regulate expression of intron-containing transcripts in response to environmental cues have been described in the budding yeast, *Saccharomyces cerevisiae* (76) and in *C. neoformans* (77). Work done on alternative splicing in *C. neoformans* supports pausing of spliceosome function (77). This fungus is intron-dense, with over 40,000 introns in its genome, and abundant alternative splicing has been observed (29). Intriguingly, the most common type of aberrant splicing is intron retention (77). Intron retention has even been shown to play a role in virulence and is regulated by environmental conditions (77). If intron retention is an adaptive mechanism for *C. neoformans* to finely tune expression levels in adverse environments, then inhibiting Prp8 intein splicing is a possible means of controlling that intron retention. Here, we propose that intron retention of transcripts is achieved through environmental sensing by the Prp8 intein, which would inhibit protein splicing and thus, cause transcripts to retain introns. The sensing ‘machinery’ of the *Cne* Prp8 intein is C1. Cysteines are powerful and reactive amino acids that endow proteins with catalytic activity, redox chemistry, and metal binding capacity (57). C1 has the full spectrum of cysteine utility, making it an excellent stress sensor.

### Nexus between protein splicing inhibition and RNA splicing

Metal sensing by the Prp8 intein *in vivo* is yet to be explored. Ideally, an inteinless strain of the native host would be needed. However, obtaining a knockout of an intein in an essential protein has proven challenging. Given these difficulties, we turned to molecular modeling of structures to help predict *in vivo* effects.

The intein-containing Prp8 precursor model generated from a solved *Schizosaccharomyces pombe* spliceosome structure revealed a snug accommodation of the intein in a cleft of Prp8 (Fig. 6A, PDB 3JB9, chain A). This insertion (site **a**) is in a linker located between the thumb domain and the endonuclease domain of the reverse transcriptase (Fig. 1B; Fig. S9). This highly conserved region of Prp8 (55%-87% identity over 113 residues) is also known as the 3’ splice site fidelity region 3.2, and is likely involved in RNA-mediated catalysis leading to intron removal (1). This site is not only at the core of the protein, but also at the catalytic center of the spliceosome. Although Prp8 likely cannot perform its function with an intein present, the structural tolerance may allow for proper folding of the intein, as well as that of Prp8. In the longer term, this flexibility gives the intein freedom to adapt to its surroundings, supporting some degree of Prp8 function in a precursor state. Mini-inteins, such as the one present in *Cne* Prp8, are not mobile and may therefore be under more selective pressure to adapt to their exteins. This is in line with work that shows partial activity of the RadA precursor with its mini-intein intact (19).

If the *Cne* Prp8 intein remains unspliced until spliceosome assembly, perhaps due to metal sensing and inhibition, it would almost certainly be disruptive. In the C complex modeled here, the intein would occupy the same location as the Cwf19 protein (Fig. 6B, Intein-protein), which is an essential spliceosome component. Cwf19 plays a central role in helping its binding partner, Prp8, fine-tune motions of the spliceosome involved in intron removal (61). If Cwf19 is unable to bind Prp8 at its cognate site, movements of the spliceosome would be altered and intron splicing disrupted. Furthermore, certain RNAs thread close to the intein (Fig. 6B, Intein-RNA). Although from our modeling there are no observable direct interactions with U2 snRNA, U2 is proximal to the intein. The binding of U2 to the intron branch-point region creates the bulged adenosine, which initiates nucleophilic attack on the 5′ splice site (78). If the intein were still present during intron removal, the critical interaction of the intron and U2 would be disturbed or changed in a way that disrupts intron splicing. If the *Cne* Prp8 intein were to sense a stressor, stay lodged in Prp8 and pause intron splicing, Prp8 precursor accumulation would undoubtedly stop the spliceosome from functioning (Fig. S10), leading to pre-mRNA accumulation, as even point mutations in Prp8 are known to do (2). Thereby, this work proposes that the *Cne* Prp8 intein is subject to modifications that would influence Prp8 function, and stall spliceosome activity. Since the observed stress-induced intein modifications are reversible, removal of the stressor would result in an immediate restoration of Prp8 and spliceosome function.

## MATERIALS AND METHODS

### Bioinformatic and phylogenetic analyses

The Prp8 intein sequences used to build the phylogenetic tree in Fig. 1A and Fig. S1A were accessed from Green et al. 2018. For comparative and phylogenetic analyses, amino acid sequences of inteins were manually trimmed to the splicing blocks (A, B, F, and G). All multiple sequence alignments of the amino acid sequences were performed using ClustalOmega with default parameters (79) and edited manually (Fig. 1; Figs. S1A and S2). Where alignments are shown shaded, black represents an identical amino acid, dark gray is a conserved amino acid, whereby the same amino acid is at the same position in a majority of the sequences, and light gray is a similar amino acid, defined as a semi-conserved amino acid substitution from the same class. Phylogenetic analysis was performed using the Neighbor-Joining (NJ) method in the MEGA7 program (80). Statistical support for the NJ tree was evaluated by Interior-branch test (number of replications, 1000) (81). The sequence logo for the B block was generated based on the multiple sequence alignment using WebLogo3 (82) (http://weblogo.threeplusone.com). The seven Prp8 intein insertions were mapped onto a model of a *Saccharomyces cerevisiae* Prp8 (Fig. S9, PDB 5GMK). All intein, Prp8, and spliceosome structures were viewed, edited, or aligned using PyMol 1.3. The 3D-BLAST protein structure search was performed by BioXGEM with default parameters (http://3d-blast.life.nctu.edu.tw).

### Bacterial strains and growth conditions

All strains used in the present study can be found in Table S1. *Escherichia coli* DH5α, MG1655(DE3), and BL21(DE3) were grown in Luria Broth (LB), unless otherwise indicated, with aeration at 250 rpm. Media contained kanamycin (50 μg/mL) or chloramphenicol (25 μg/mL) where appropriate. Plasmids were transformed into cells by electroporation using a Bio-Rad Gene Pulser and recovered for 1 h at 37°C in SOC medium (0.5% yeast extract, 2% tryptone, 10 mM NaCl, 2.5 mM KCl, 10 mM MgCl_2_, 10 mM MgSO_4_, and 20 mM glucose). Transformants were selected by plating on LB agar with the appropriate antibiotic and incubated at 37°C overnight.

### Construction of plasmids

All plasmids used in the present study can be found in Table S2 and all oligonucleotides, synthesized by Integrated DNA Technologies (IDT), are in Table S3. Plasmid DNA was prepared using E.Z.N.A. Plasmid Mini Kit (Omega). DNA was visualized in 1% agarose gels using EZ-Vision DNA Dye (Amresco). PCR fragments were amplified using CloneAmp HiFi PCR Premix (Clontech) from genomic DNA of *C. neoformans* var. grubii H99 or *C. gattii* NIH444 (Dr. Sudha Chaturvedi, New York State Department of Health), *A. fumigatus* AF293 (Dr. Robert J. Cramer, Darmouth College), *B. dendrobatitidis* JEL423 (Dr. Timothy James, University of Michigan), or *H. capsulatum* G186A (Dr. Chad Rappleye, Ohio State University). For insertion into the MIG construct, the inserts included 5 native N- and C-extein residues flanking the intein. For insertion into the overexpression vector, pET47b, the intein alone with 3 native N-exteins was PCR amplified. Digested backbone was gel purified using Zymoclean Gel DNA Recovery Kit (Zymo Research). Restriction enzymes (NEB), T4 ligase (NEB), and In-Fusion HD Cloning Plus Kit (Clontech) were all used per manufacturer protocol. Mutagenesis was performed using the QuikChange Lightning Site-Directed Mutagenesis Kit (Agilent) for single amino acid mutations or the QuikChange Lightning Multi Site-Directed Mutagenesis Kit (Agilent) for multiple amino acid mutations. For the A-1V mutation, primers were designed to randomly mutate the A-1 to all other possible codons using a degenerate primer with NNS at the mutated location. All clones were verified by sequencing (EtonBio).

### MIG splicing assays

MIG Prp8 (WT, A-1V, and derived C61 mutants) was transformed by electroporation into MG1655(DE3). The cells were subcultured 1:100 from an overnight culture into fresh LB medium and grown at 37°C with 250 rpm shaking to an OD_600_ of 0.5. Cells were induced with 0.5 mM IPTG for 1 h at 30°C and pelleted by spinning for 10 min at 4,000 rpm. The pellets were lysed immediately using tip sonication (20 sec on/30 sec off at 30% amplitude for 1 min total) in 50 mM Tris, pH 8.0, and 10% glycerol or stored at −80°C until lysis. For any ROS/RNS or metal treatment, the indicated compound was added to cells at the desired concentration prior to incubation at 30°C for the specified time. EDTA was added to a final concentration of 10 mM and TCEP to a final concentration of 40 mM. Upon completion of the assay or time point, the lysate was frozen at −80°C. To visualize MIG splicing assay results, samples were separated under non-reducing conditions on Novex WedgeWell 12% Tris-Glycine gels (Invitrogen) using loading dye lacking β-mercaptoethanol and visualized using a Typhoon 9400 scanner (GE Healthcare) with excitation at 488 nm and emission at 526 nm. Quantitation and analysis were done using ImageJ and GraphPad Prism (v7.02).

### Prp8 intein purification

For isothermal titration calorimetry and mass spectrometry, the *Cne* Prp8 intein from *Cryptococcus neoformans* var. *grubii* H99 was amplified with 3 native N-extein residues (EKA) and cloned into pET47b in front of an N-terminal His_6_-tag and an HRV 3C protease site. For crystallization, the *Cne* Prp8 intein with 2 native N-extein residues (KA) and the native C-extein S+1 was amplified and cloned into pET28a with a C-terminal His_6_-tag using a megaprimer approached as described previously (83).

The pET47b (or pET28a) *Cne* Prp8 intein construct was transformed by electroporation into BL21(DE3) cells. The cells were subcultured 1:100 from an overnight culture into fresh LB medium and grown to an OD_600_ of 0.6. Cells were induced with 0.5 mM IPTG and grown with shaking at 250 rpm overnight at 16°C. The following morning, cells were harvested by centrifugation at 4,000 rpm for 10 minutes. Pellets were frozen at −80°C until ready for lysis. Tip sonication was performed (30 sec on/59 sec off at 30% amplitude for 4 min total) in buffer containing 20 mM Tris, pH 7.8, 500 mM NaCl, 25 mM imidazole, and 5% glycerol. Whole cell lysate was centrifuged at 20,000 × *g* for 20 min to separate the soluble fraction, which was loaded onto a nickel affinity column equilibrated with the lysis buffer. Washes were carried out using buffer containing 20 mM Tris, pH 7.8, 500 mM NaCl, 75 mM imidazole, and 5% glycerol and elution buffer with 20 mM Tris, pH 7.8, 500 mM NaCl, 250 mM imidazole, and 5% glycerol. Purified fractions of the *Cne* Prp8 intein were checked by separation on SDS-PAGE and the cleanest elution samples were pooled. For the pET47b construct, the His_6_-tag on the *Cne* Prp8 intein was removed through digestion with HRV 3C protease according to the manufacturer’s protocol. The cleaved *Cne* Prp8 intein reaction was passed back over a nickel affinity column and the flow-through was collected to ensure no His_6_-tagged *Cne* Prp8 intein or HRV 3C protease was in the sample. For analysis by ITC, the flow-through intein was exchanged into 50 mM C_2_H_3_NaO_2_ (pH 7.0), 100 mM NaCl using a HiPrep 26/10 desalting column or a dialysis cassette. For mass spectrometry, the flow-through intein was used directly for metal treatments and then further purified by liquid chromatography (LC) prior to spraying on the instrument. For the pET28a construct, the imidazole-eluted fractions were concentrated and subjected to size exclusion chromatography by a gel filtrations 16/60 Superdex column (GE Healthcare). For crystallization, the purified *Cne* Prp8 intein was concentrated to 9.5 mg/mL in a buffer composed of 25 mM HEPES, pH 7.5, and 150 mM NaCl.

### Mass spectrometry of Prp8 intein

Purified *Cne* Prp8 intein was reduced with 40 mM TCEP and exchanged into deoxygenated exchange buffer (20 mM Tris pH 7.5, 200 mM NaCl) using 7K MWCO Zeba spin desalting columns (Thermo) to remove TCEP. The protein concentration was measured and then treated with 10X of CuSO_4_ and incubated at 30°C for 1 h. Following treatment, the purified intein was denatured with 6 M urea at 37°C for 30 min. The urea concentration was diluted down to less than 0.8 M with 50 mM Tris, pH 7.6, 1 mM CaCl_2_. Trypsin digest of the intein was performed by adding activated trypsin (Promega) to a final ratio of 1:20 and incubating overnight at 37°C. The oxidation of *Cne* Prp8 intein cysteines after treatment was analyzed by multiple reaction monitoring-initiated detection and sequencing (MIDAS) as described (84). The trypsin-digested mixture was acidified followed by LC-MS/MS analysis. LC-MS/MS analysis was performed on a microflow LC-MS/MS system configured with a 3-pumping Micromass/Waters CapLC™ system with an autosampler, a stream select module configured for precolumn plus analytical capillary column, and a QTRAP 6500 (ABSCIEX) mass spectrometer fitted with Turbo V microflow source, operated under Analyst 1.63 control. Injected samples were first trapped and desalted isocratically on an LC-Packings PepMap™ C18 μ-Precolumn™ Cartridge (5 μm, 500 μm I.D. × 20 mm; Dionex, Sunnyvale, CA, USA) for 7 min with 0.1% formic acid delivered by the auxiliary pump at 40 μL/min after which the peptides were eluted from the precolumn and separated on an analytical C18 capillary column (15 cm × 500 μm i.d., packed with 5 μm, Jupiter 300 C18 particles, Phenomenex, CA, USA) connected inline to the mass spectrometer, at μL/min using a 50 min gradient of 5% to 80% acetonitrile in 0.1% formic acid. The oxidized peptide identification was conducted through multiple reaction monitoring (MRM) triggered enhanced product ion (EPI) scan using information dependent acquisition (IDA). The utilization of chromatographic separation, MRM transitions, and EPI scan allows accurate peptide identification and confirmation. The two MRM transitions including m/z 404.19 > 532.22 and m/z 786.04 > 895.41 for C[Oxi]LQNGTR.+2b5 and THEGLEDLVC[Oxi]THNHILSMYK.+3b8 were used to trigger the EPI experiment respectively. The instrument was operated in a positive ion mode with a turbo V ion drive electrospray source. The parameters for the operation were as follows: curtain gas, 20 psi; heated nebulizer temperature 180°C, ion spray voltage, 5500 V; gas1, 18 psi; gas 2, 15 psi, de-clustering potential, 65 V, EP, 10 V and CAD gas, high.

### Isothermal titration calorimetry of Prp8 intein

ITC measurements were carried out on a TA Instruments Nano ITC (TA Instruments, Inc., New Castle, DE). Aqueous solutions of metal titrants (CuSO_4_ or ZnSO_4_) were prepared to be 0.3-30-fold higher than the concentration of the *Cne* Prp8 intein, in the range of 0.05-5.0 mM. The titrant and *Cne* Prp8 intein samples (wild-type and mutants) were degassed before each titration. The purified *Cne* Prp8 intein was concentrated from 10 μM to 16 μM in 300 μL and were placed in a 2.5 mL reaction cell, and the reference cell was filled with 300 μL deionized water. All titrations were carried out at 37°C. After baseline equilibration, successive injections of an indicated titrant were made into the reaction cell in 2.5 μL increments at 400 s intervals with stirring at 250-350 rpm to ensure an equilibrium was achieved for a return to baseline. The resulting heats of reaction were measured over 20 consecutive injections. Buffer control experiments [50 mM C_2_H_3_NaO_2_ (pH 7.0), 100 mM NaCl, ± 10 mM tris(2-carboxyethyl)phosphine] to determine the heats of titrant dilution were carried out by making identical injections in the absence of the *Cne* Prp8 intein. The net reaction heat was obtained by subtracting the heats of dilution from the corresponding total heat of reaction. The titration data were deconvoluted based on best-fit binding models containing either one or three sets of interacting binding sites, using a nonlinear least-square algorithm through the NanoAnalyze software. The binding enthalpy change (ΔH), dissociation constant (K_d_), and binding stoichiometry (n) were permitted to vary during the least-square minimization process, and taken as best-fit values.

### Crystallization, structure determination, and refinement of Prp8 intein

Initial crystallization conditions were obtained by screening the Hampton crystallization screens (I, II, and Research Index HT), using the hanging-drop vapor diffusion method. Upon optimization, large crystals were grown by mixing 1 μL of *Cne* Prp8 intein and 1 μL of reservoir solution containing 22-28% PEG4,000, 0.1 M sodium acetate, pH 4.2, 0.2 M ammonium acetate. The *Cne* Prp8 intein crystallizes in space group *P1* with six intein molecules per asymmetric unit. Prior to data collection, all crystals were transferred to a cryo-protectant solution containing crystallization buffer supplemented with 20% glycerol. The crystals were flash-cooled directly in liquid nitrogen. Diffraction data for the co-crystals were collected at 100 K using a Pilatus detector at the BL9-2 beamline of the Stanford Synchrotron Radiation Laboratory. Data were processed, scaled, and reduced using the programs HKL2000 (85) and Phenix suite (86). The structure of the *Cne* Prp8 intein was determined by molecular replacement, with the crystal structure of the *Cryptococcus gattii* Prp8 intein (Li et al., in prep) as a search model using the PHENIX program suite. Structure refinement was carried out using the Phenix program suite (Table S4).

### Modeling of zinc binding

The models for zinc ion binding were constructed using the *Cne* Prp8 intein crystal structure conformation (chain A) in the asymmetric unit and a structure deposited in the PDB of human ubiquitin ligase (87), which has a zinc atom bound to two cysteines and two histidine side chains (PDB 5TDA). This ubiquitin ligase structure was chosen because of its high resolution (0.79 Å), and the occurrence of the four side chains present in the *Cne* Prp8 intein coordinating a zinc ion. All the optimizations were performed using the program CHARMM (version c35b3) (88, 89) and the CHARMM36 force field for proteins (90). The best-fit orientations of a deprotonated C1, a deprotonated C61, a neutral H63 side chain, and a neutral H65 side chain of each Prp8 intein crystal structure conformation onto the corresponding C112, C115, H133, and H136 side chains of the human ubiquitin ligase were first obtained using a Kabsch least-squared optimization (91). The coordinates of the zinc ion were then transferred from the ubiquitin ligase structure, and subsequently held fixed. All protein non-hydrogen atoms, except those in C1, C61, H63, and H65, were harmonically restrained with a force constant of 1 kcal/mol/Å^2^ to prevent unrelated structural changes in the intein. Root Mean Square deviation (RMSD) restraints were then applied to the C1, C61, H63, and H65 residues to gradually change their conformation to a zinc-coordinating one, as observed in the ubiquitin ligase structure. The RMSD restraints had a force constant of 10000 kcal/mol/Å^2^, were applied only to non-hydrogen atoms, and the restraint minimum was gradually changed in 0.5 Å decrements from an initial RMSD of 4.5 Å to a final RMSD of 0 Å. For each restraint minimum value, the following optimization protocol was followed: a Steepest Descent (SD) (92) minimization for 5000 steps with an energy change tolerance criterion of 0.001 kcal/mol, 1000 steps of Langevin dynamics at a temperature of 150 Kelvin and a friction coefficient of 60.0 ps^−1^, another 5000 SD minimization steps with an energy change tolerance criterion of 0.001 kcal/mol. Each model thus obtained was further minimized using 5000 SD steps with an energy change tolerance criterion of 0.001 kcal/mol, the harmonic restraints on the other protein atoms were removed, and a further 5000 SD steps of minimization with an energy change tolerance criterion of 0.001 kcal/mol were carried out to get the final model.

### Precursor modeling

The Prp8 protein structure (chain A) in the 3.6 Å cryo-EM structure of the *Schizosaccharomyces pombe* (*Spo*) spliceosome (12) (PDB 3JB9) was used as a structural template for constructing a homology model of the *Cne* Prp8 exteins. A pairwise Needleman-Wunsch (93) sequence alignment using EMBOSS Needle (94) shows 70.6% sequence identity and 83.1% sequence similarity between the full *Spo* and *Cne* extein sequences. The *Cne* Prp8 extein homology model was constructed using MODELLER (95) using the DOPE (96) and GA341 (97) energy functions to identify the best model. The *Spo* Prp8 extein template has longer missing sections at the N- and C-termini, as well as some smaller gaps in between (the missing residues are 1-46, 303-313, 1533-1538, 1781-1783, and 2031-2363). The 171 residue *Cne* Prp8 intein sequence is an insert between residues 1530 and 1531 in the extein, which suggests only a relatively short 5-residue gap (*Spo* Prp8 residues 1533-1538) as a factor limiting modeling accuracy of the precursor state. The *Cne* Prp8 extein homology model was then combined with the *Cne* Prp8 intein crystal structure to generate a model for the full *Cne* Prp8 precursor. The orientation of the intein with respect to the extein was manually adjusted using VMD software (98) to avoid any steric overlap and keep the intein and extein ends that are joined by peptide bonds sufficiently close together. This combined intein-extein structure was then used as a structural template to generate a continuous *Cne* Prp8 precursor homology model using MODELLER. Further minimization on this precursor homology model was performed using the program CHARMM, version c35b3 (88, 89) with the CHARMM36 force field for proteins (90). All atoms not within the precursor amino acid sequence range encompassing the intein and its neighboring extein regions (residues 1520-1720) were initially held fixed, and a low-temperature (150 K) optimization protocol was used to improve the homology model. This protocol included 5 iterations of the following steps: (a) 5000 steps of Steepest Descent (SD) minimization followed by 5000 steps of Adapted-Basis Newton-Raphson (ABNR) minimization, each with an energy change tolerance of 0.001 kcal/mol, (b) 1000 steps of Langevin dynamics at a temperature of 150 Kelvin and a friction coefficient of 5.0 ps^−1^, (c) another 5000 steps of SD and 5000 steps of ABNR minimization. All non-hydrogen atoms were then restrained using harmonic restraints with a force constant of 1.0 kcal/mol/Å^2^ and SHAKE constraints (99) were applied on all hydrogen atoms, and 5000 steps of SD and 5000 steps of ABNR minimization was performed to obtain the final *Cne* Prp8 precursor model.

## ACKNOWLEDGEMENTS

The authors acknowledge Dr. Qishan Lin (UAlbany) for help with mass spectrometry experiments. Fungal genomic DNA was donated by Dr. Sudha Chaturvedi (Wadsworth Center), Dr. Robert Cramer (Dartmouth), Dr. Timothy James (University of Michigan), and Dr. Chad Rappleye (Ohio State). We also wish to thank Dr. Tzanko I. Doukov at the Stanford Synchrotron Radiation Lightsource (SSRL) for help in X-ray diffraction data collection. Use of SSRL, SLAC National Accelerator Laboratory, is supported by the U.S. Department of Energy (DOE) under Contract No. DE-AC02-76SF00515. The SSRL Structural Molecular Biology Program is supported by the DOE and by the National Institutes of Health (NIH), including P41GM103393. This work was supported by the NIH grants GM39422 and GM44844 to M.B.

The atomic coordinates and structure factors (PDB 6MX6) have been deposited in the Protein Data Bank, Research Collaboratory for Structural Bioinformatics, Rutgers University, New Brunswick, NJ (http://www.rcsb.org/).

## SUPPLEMENTAL INFORMATION

**Figure S1.**
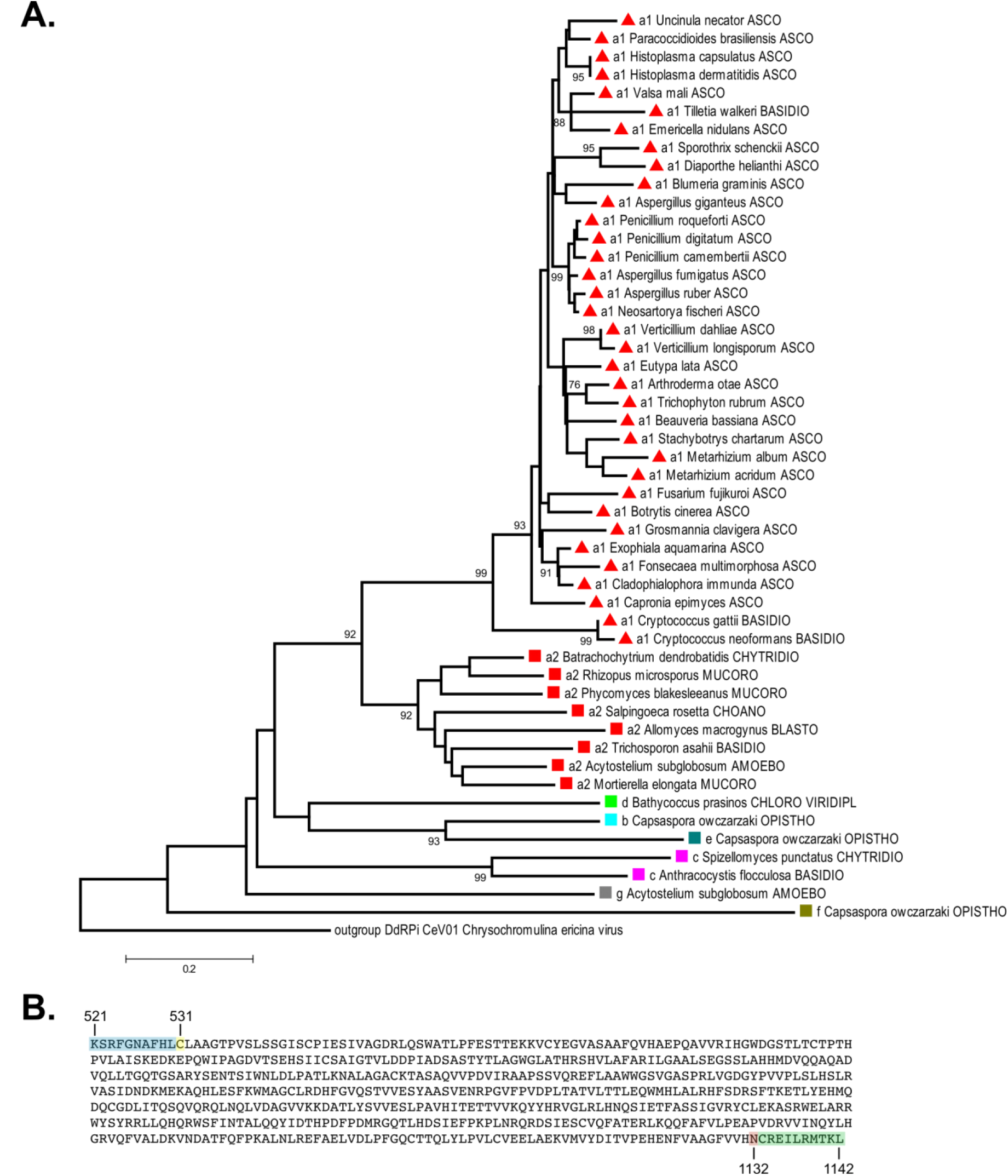
Distribution of Prp8 inteins and novel insertion site **g. A.** A phylogenetic tree of Prp8 inteins was reconstructed based on an amino acid multiple sequence alignment of the splicing blocks (A, B, F, G) using the Neighbor-Joining algorithm and an Interior-branch test with 1000 replicates. Fifty representatives covering Prp8 intein diversity were selected and the full name of each intein-containing organism is listed. Colored symbols represent the insertion site and correspond to colors in Fig. 1A. Letters (a1, a2, b, c, d, e, f, g) represent each of the seven unique insertion sites. Phylum abbreviations are as follows: Amoebo – Amoebozoa; Asco – Ascomycota; Basidio – Basidiomycota; Blasto – Blastocladiomycota; Choano – Choanoflagellida; Chloro Viridipl – Chlorophyta Viridiplantae; Chytridio – Chytridiomycota; Mucoro – Mucoromycota; Opistho – Opisthokonta. **B.** Novel Prp8 insertion site **g**. In the amoeba *Acytostelium subglobosum* (*Asu*), an intein was identified at a new site in Prp8, here termed **g**. This is the seventh site in which a Prp8 intein has been found. The full site **g** intein sequence is shown, plus 10 flanking N-extein (blue) and C-extein (green) amino acids. The *Asu* C1 (yellow) and terminal asparagine (red) are highlighted. Residue numbering corresponds to the *Asu* Prp8 exteins. Accession number: XP_012753295.

**Figure S2.**
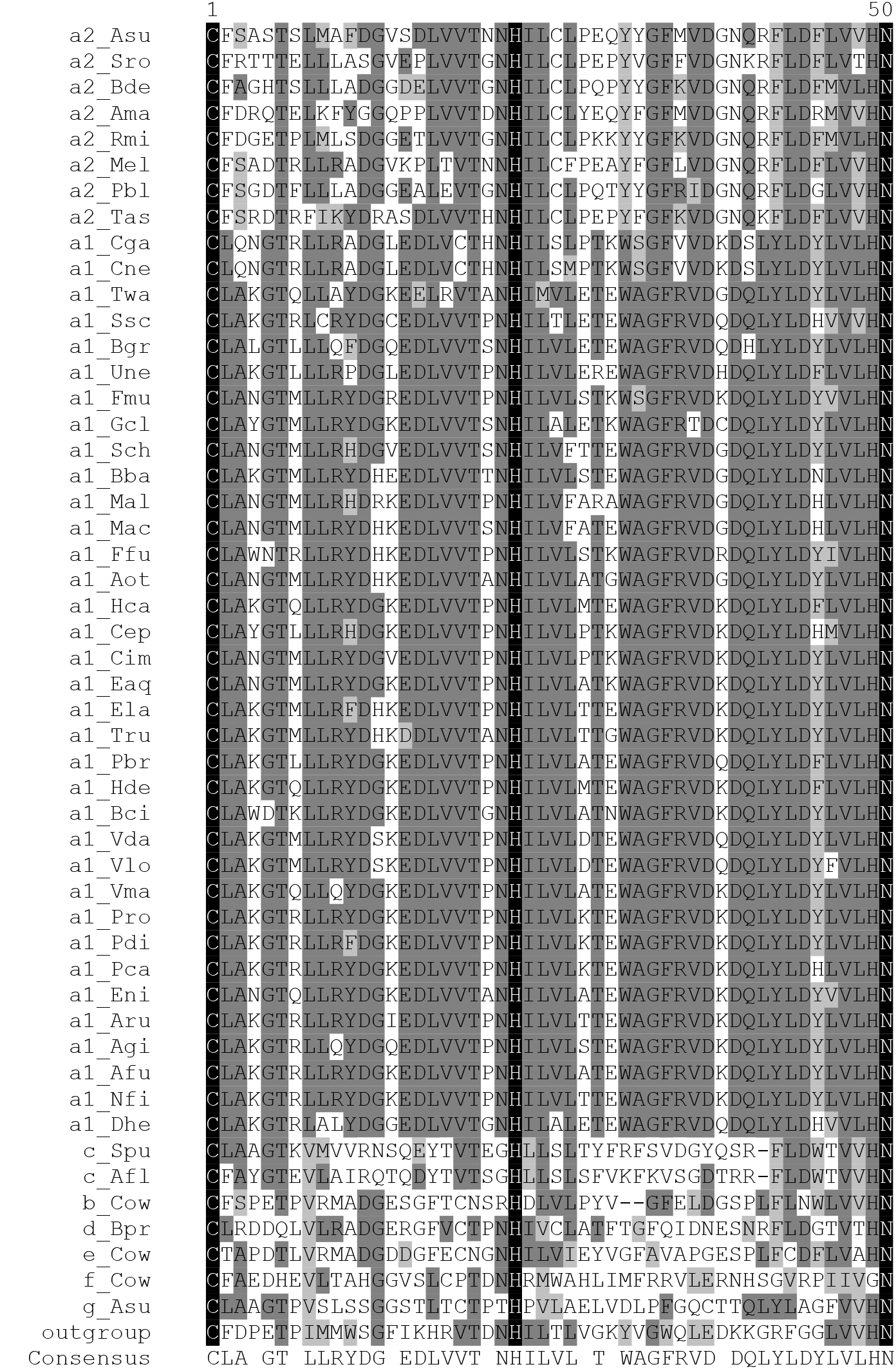
Amino acid multiple sequence alignment of Prp8 inteins utilized for phylogenetic analysis. Comparative analysis of amino acid residues found in blocks A, B, F, and G from the selected 50 representative Prp8 inteins, shown with abbreviated species names (full names in Fig. S1). Letters (a1, a2, b, c, d, e, f, g) represent each of the seven unique insertion sites. Shading is as follows: black – identical amino acid, dark gray – conserved amino acid, light gray – similar amino acid substitution.

**Figure S3.**
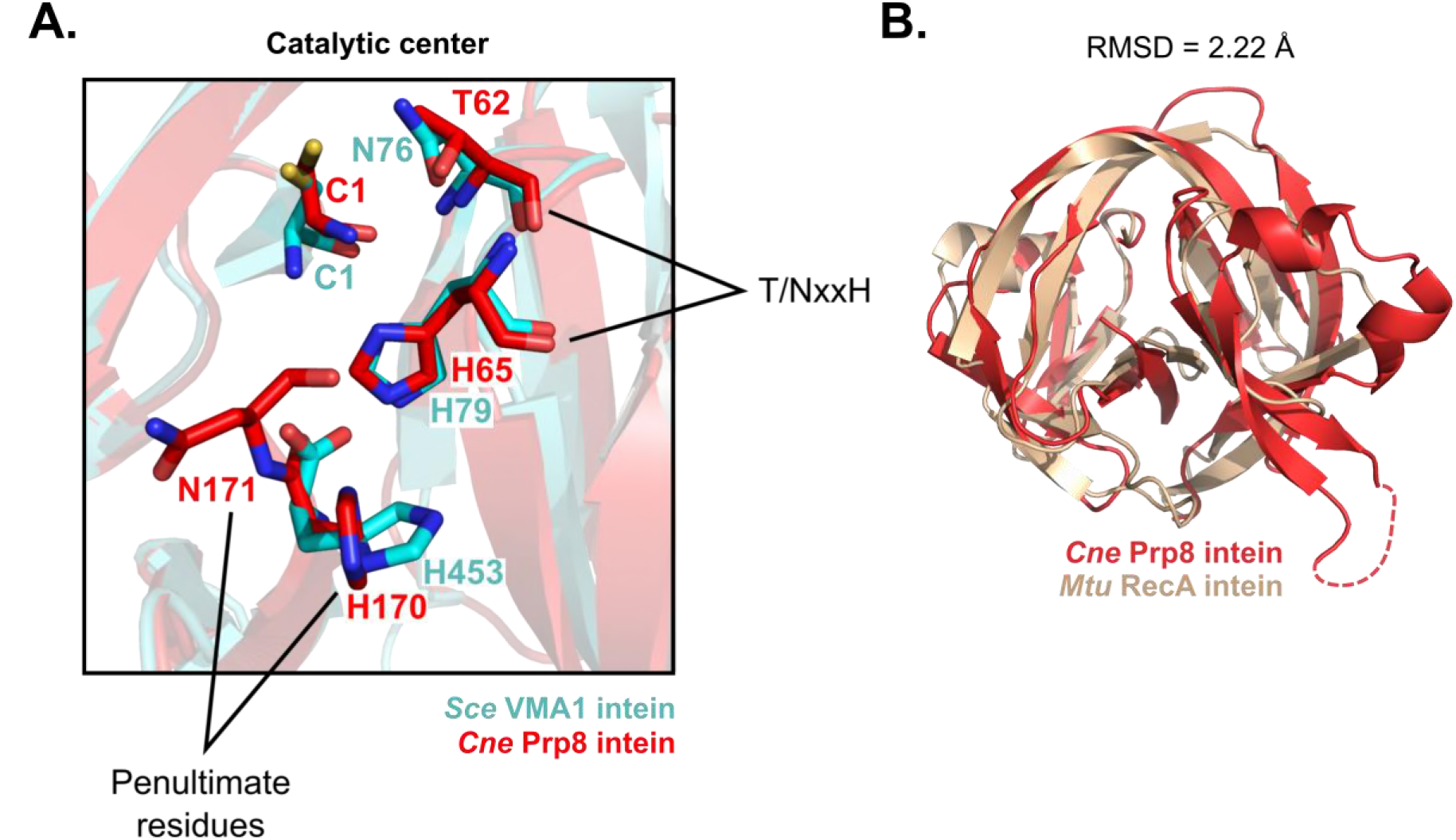
Overlays of the *Cne* Prp8 intein with other inteins. **A.** Overlay of the *Saccharomyces cerevisiae* (*Sce*) VMA1 intein and *Cne* Prp8 intein active sites. The *Sce* VMA1 intein (cyan, PDB 1GPP) was overlaid with the *Cne* Prp8 intein (red). The active site residues, crucial to protein splicing, are shown as sticks and labeled. A majority of these conserved residues overlap exactly, such as the catalytic C1, and the B block TxxH motif. The *Sce* VMA1 intein uses an asparagine (N76) rather than threonine in the TxxH motif, but the positioning is similar to the threonine (T62) of the *Cne* Prp8 intein. The penultimate histidines (H170 and H453) are in comparable positions except for the side chains, whose chi angles are different by 45°. The *Sce* VMA1 intein was not solved with the terminal asparagine. **B.** Structural comparison of bacterial *Mycobacterium tuberculosis* (*Mtu*) RecA intein and fungal *Cne* Prp8 intein. Overlay of the *Mtu* RecA intein (brown, PDB 2IMZ) and the *Cne* Prp8 intein (red) reveals structural similarities in major intein features, such as the anti-parallel β-sheet folding, that contribute to the horseshoe shape. The Hint domain, comprised of splicing blocks A, B, F, and G, are generally aligned between the two inteins. The structures deviate at sequences between blocks B and F, where the *Cne* Prp8 intein encoded a linker or endonuclease domain. The two structures have an RMSD value of 2.22 Å.

**Figure S4.**
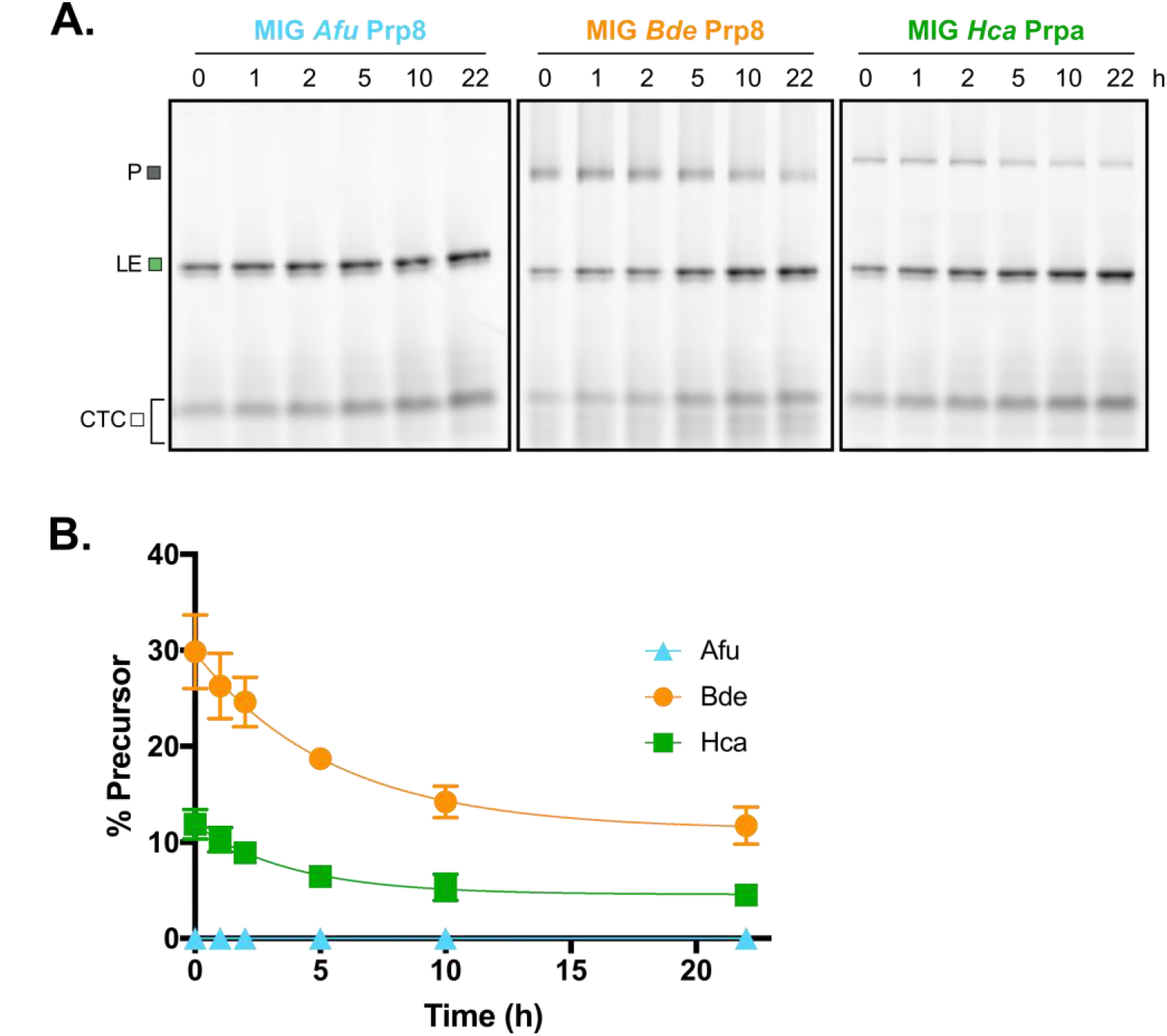
Splicing of Prp8-**a** inteins from other fungal pathogens in MIG. **A.** Diverse Prp8 intein splicing patterns. Several other Prp8 inteins from human fungal pathogens *Aspergillus fumigatus* (*Afu*), *Batrachochytrium dendrobatidis* (*Bde*), and *Histoplasma capsulatum* (*Hca*) were cloned into MIG. Splicing was observed over time by the loss of precursor (P) and increase in ligated exteins (LE), or simply by the presence of ligated exteins (for *Afu*). The gel shows that not all Prp8 inteins splice similarly, despite being placed in an identical extein context. **B**. Precursor amounts vary greatly. A quantitation of precursor (P) at each time-point shows that these Prp8 inteins are active, but splice at variable rates. The *Afu* Prp8 intein is almost entirely spliced at the start of the assay (0 h), whereas *Bde* has 31% precursor at 0 h and *Hca* has 14% precursor at 0 h. The variable rates were determined by calculating the loss of precursor over time for MIG *Bde* Prp8 and MIG *Hca* Prp8, and are 3.6 × 10^−2^ per min and 1.9 × 10^−2^ per min, respectively. This suggests intein-mediated control of protein splicing. Data are representative of three biological replicates and mean standard deviations are shown. Trend lines are fit to show the decay curve.

**Figure S5.**
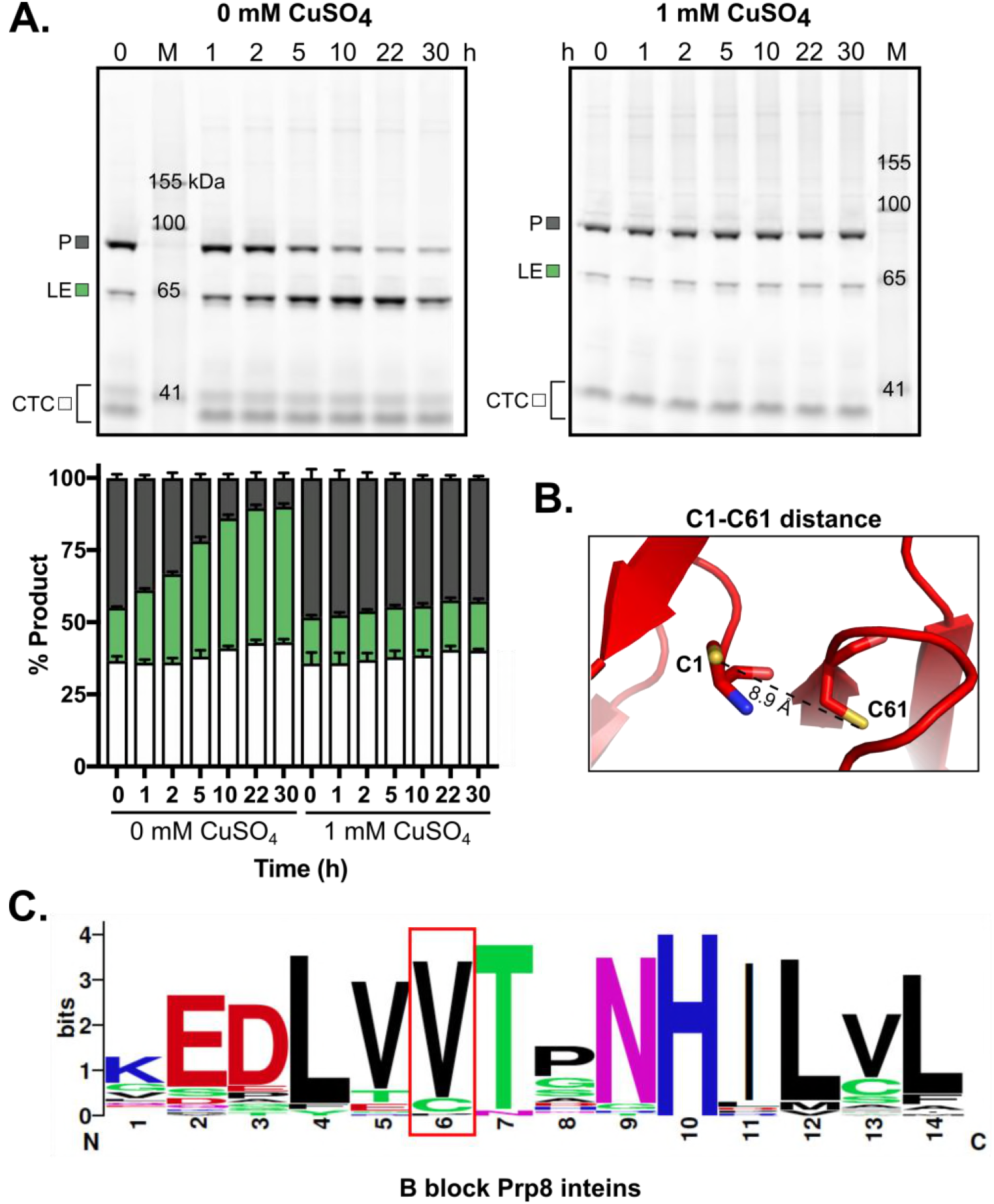
MIG Prp8 A-1V copper inhibition and cysteine analysis. **A.** Copper treatment causes inhibition. Induced MIG Prp8 A-1V cells were lysed and treated with 0 or 1 mM CuSO_4_. The lysates were incubated for the indicated time at 30°C and then frozen. Samples were separated on SDS-PAGE and scanned for GFP fluorescence. In the absence of copper, MIG Prp8 A-1V spliced well over 30 h, converting precursor (P) into ligated exteins (LE). There was little to no conversion of P to LE over time with copper was addition. The inhibition was still observed at 30 h after copper addition. Quantitation is shown below in a stacked plot. Data are representative of three biological replicates and mean standard deviations are shown. **B.** Relative position of two cysteines. There are only two cysteines present in the *Cne* Prp8 intein. Using the solved structure, a measurement of the distance between C1 and C61 (shown as sticks) was calculated to be 8.9 Å apart. **C.** Valine is the preferred residue at position 61. A sequence logo was constructed of the B block from 50 representative Prp8 inteins. This shows absolute conservation of the histidine (position 10) and a strong preference for threonine (position 7) in the TxxH motif. However, the B block cysteine (position 6, red box) is not highly conserved across Prp8 inteins and most encode valine at this site.

**Figure S6.**
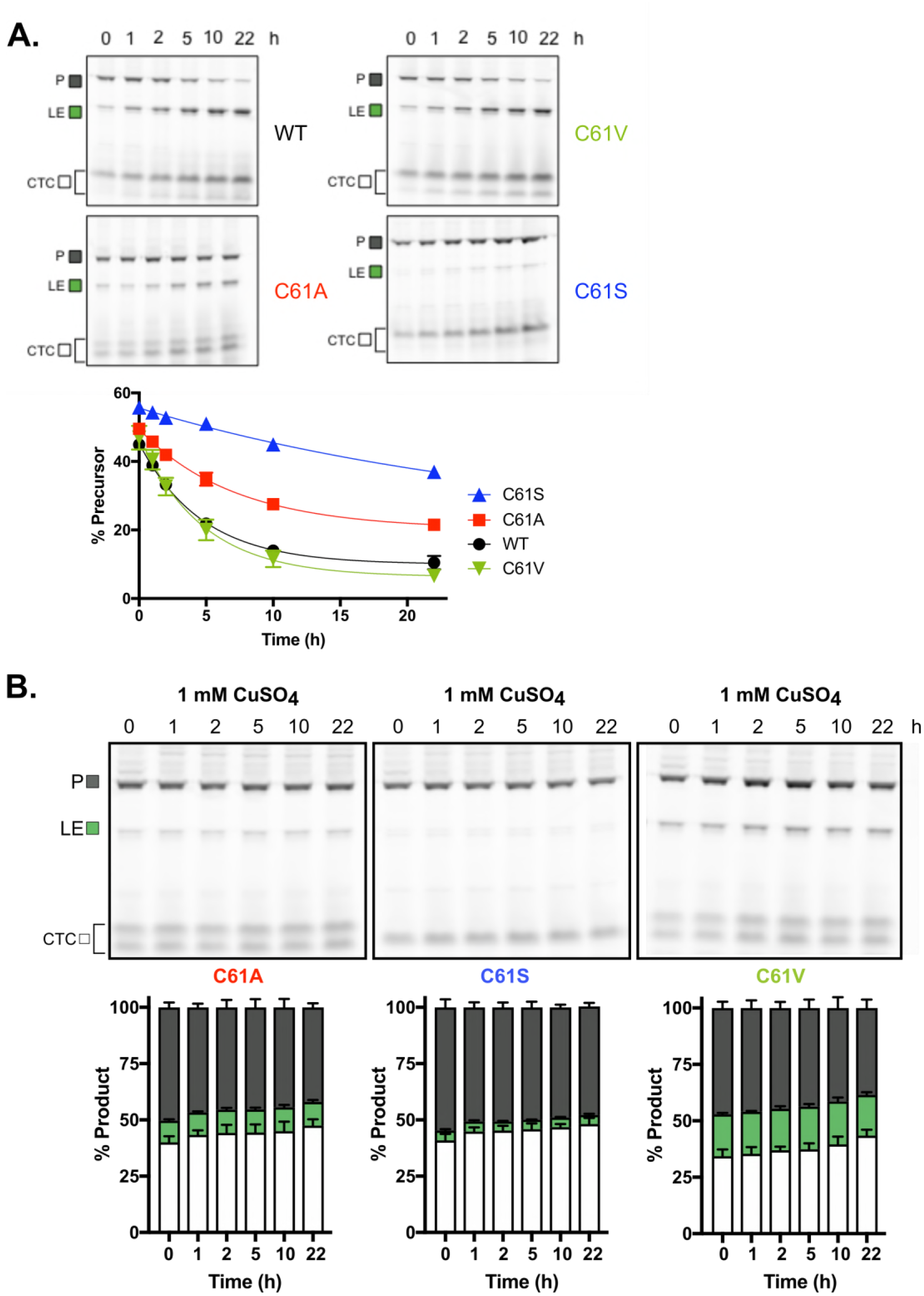
Copper inhibition of MIG Prp8 A-1V C61 mutants. **A.** Mutations to C61 in MIG Prp8 A-1V affect splicing rates. The B block C61 was mutated to valine (C61V), alanine (C61A), and serine (C61S) and splicing observed over time in MIG. Splicing rates were determined by calculating the loss of precursor over time and are as follows: WT – 9.2 × 10^−2^ per min, C61V – 8.6 × 10^−2^ per min, C61A – 4.7 × 10^−2^ per min, and C61S – 1.6 × 10^−2^ per min. The C61V mutant splices similarly to WT, whereas C61A and C61S are slower. A quantitation is shown to the right with the amount of precursor (P) at each time-point. Data are representative of three biological replicates and mean standard deviations are shown. Trend lines are fit to show the decay curve. **B.** MIG Prp8 A-1V B block cysteine mutants are inhibited by copper. To test if copper inhibition was caused by C1 oxidation, C61 mutants were treated with CuSO_4_. After induction of MIG, the cells were lysed and 1 mM CuSO_4_ was added. The lysates were incubated at 30°C and aliquots were collected at the indicated time. Samples were run on SDS-PAGE and scanned for GFP fluorescence. None of the C61 mutants show an increase in ligated exteins (LE) over time, with little loss of precursor (P). This indicates that at least C1 oxidation by copper is sufficient to cause the observed splicing inhibition, and that disulfide bonds are not involved. Quantitation is shown below in a stacked plot. Data are representative of three biological replicates and mean standard deviations are shown.

**Figure S7.**
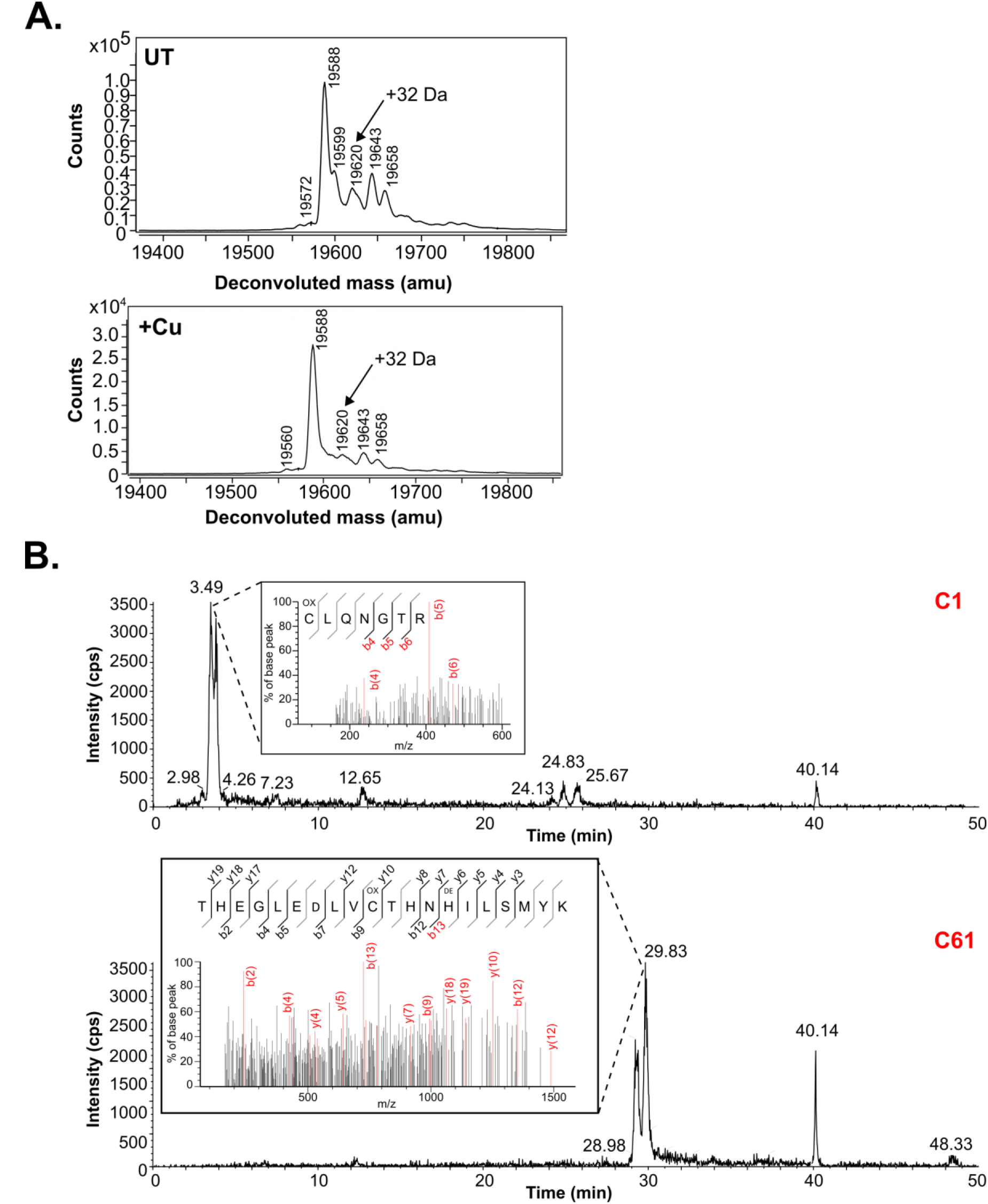
Mass spectrometry of cysteine modifications. **A.** Intact *Cne* Prp8 intein shows small mass shift. Purified *Cne* Prp8 intein was untreated or treated with excess copper and separated and analyzed using LC-MS. The peaks were deconvoluted and the expected mass of the Prp8 intein, 19588 Da, is seen as the largest peak. A small, 32 Da shift (19620 Da) was visible with both no treatment and copper treatment only (arrow). This suggests highly reactive cysteines that are modifiable by atmospheric oxygen alone. **B.** C1 and C61 are oxidized with copper treatment. Trypsin-digested fragments of copper-treated *Cne* Prp8 intein were separated and sprayed using LC-MS/MS (insets). Peptides (red peaks) containing C1 or C61 were detected and further analyzed using multiple reaction monitoring-initiated detection and sequencing (MIDAS) to confirm the identity and location of oxidation. The chromatogram shows elution time for both cysteines consistent with a single additional oxygen, or a sulfenic acid modification.

**Figure S8.**
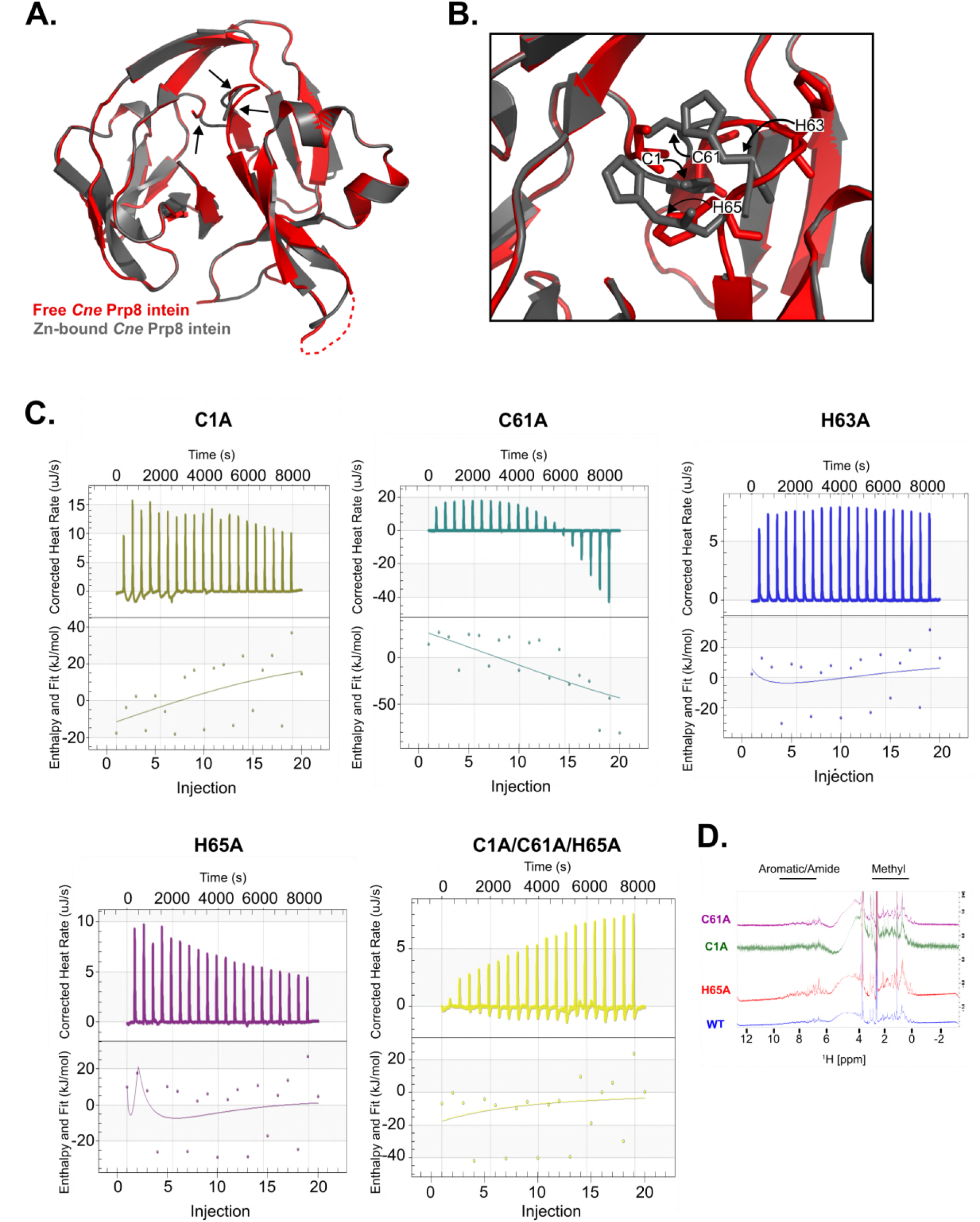
Putative zinc binding site. **A.** Overlay of the free *Cne* Prp8 intein with zinc-bound intein model. The free *Cne* Prp8 intein (red) is overlaid with the optimized zinc-binding model (gray). Minor conformational changes of the intein are required for the pocket to form, indicated by arrows, with an RMSD of 0.104 Å. **B.** Zoom into zinc coordination residues. The residues suspected to form a binding pocket (C1, C61, H63, and H65) are shown as sticks in the free intein structure (red) or zinc-bound intein model (gray). Rotational freedom of these amino acid side chains allows movement of the residues to coordinate the zinc ion. Arrows indicate direction of movement from free intein to zinc-bound structure. **C.** Using isothermal titration calorimetry (ITC), purified 16 μM Prp8 intein with various binding site mutations was titrated with 0.5 mM ZnSO_4_ over 20 injections at 37°C and pH 7.0 on a Nano ITC. Changes in heat after incremental ZnSO_4_ injections are shown in the top panel and are corrected by subtracting the heat of ZnSO_4_ titrated into buffer alone. The bottom plot shows integrated heat per mole of ZnSO_4_ as a function of the molar ratio of ZnSO_4_ to Prp8 intein. Binding was not detected in a biologically significant range for all mutants compared to the wild-type, suggesting that one, two, or all three residues play a role in zinc binding to the *Cne* Prp8 intein (see Table 1). Although the ITC data integration and binding model affirmation occasionally produced high error values, experiments were conducted in triplicate, and persistently completed highly reproducible enthalpic transitions. Repeated, corresponding transition levels along the binding isotherms at stoichiometric equivalent points consistently approached unity values despite these errors. Representative data is shown. **D.** 1D proton NMR spectrum of WT *Cne* Prp8 intein is shown (blue) and overlaid with the same spectra of three representative mutants (C1A, C61A, and H65A). The Aromatic/Amide amino acid region is similar across all proteins, suggesting that the mutants fold and are structurally sound.

**Figure S9.**
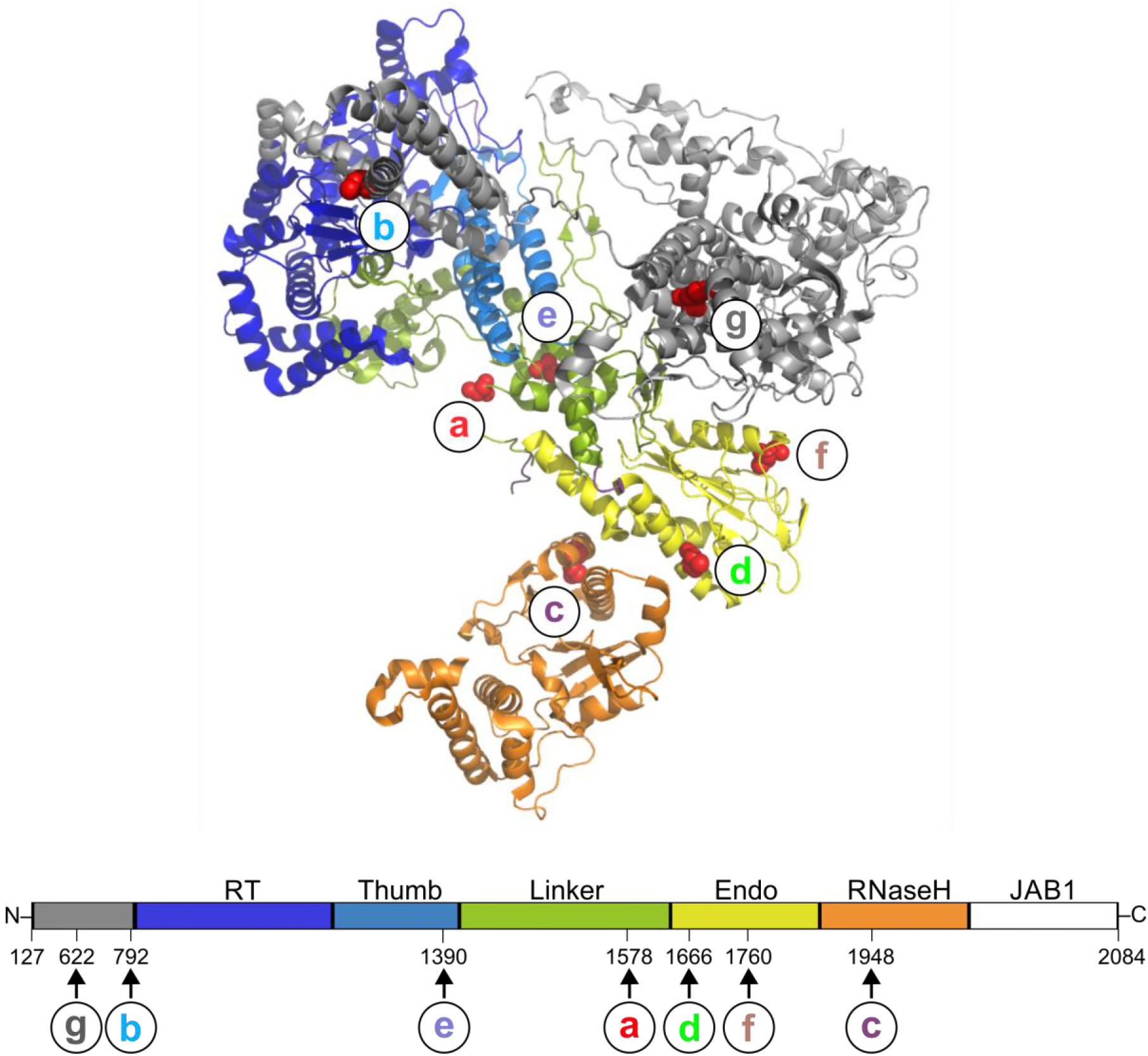
Mapping of Prp8 intein insertion sites to Prp8 extein domains. The seven unique insertion sites (**a-g**) were mapped to a solved structure of Prp8 from a *Saccharomyces cerevisiae* C complex spliceosome (PDB 5GMK, chain A from Wan et. al, 2016) by locating the +1 residue. This Prp8 structure was used because the insertion sites are all resolved. The +1 residues are shown as red spheres and labeled **a-g**. Most Prp8 inteins localize close to the active center of Prp8. Some insertions are in the N-terminal domain, which provides structural integrity to the spliceosome. A corresponding line diagram of Prp8 exteins shows the domains of the host protein from amino acid residues 127 to 2084 with arrows indicating the site of intein insertion with the residue number and insertion site letter. The domains are as follows: N-terminal domain – gray, RT Palm/Finger – dark blue, Thumb/X – light blue, linker – green, endonuclease – yellow, and RNase H-like – orange.

**Figure S10.**
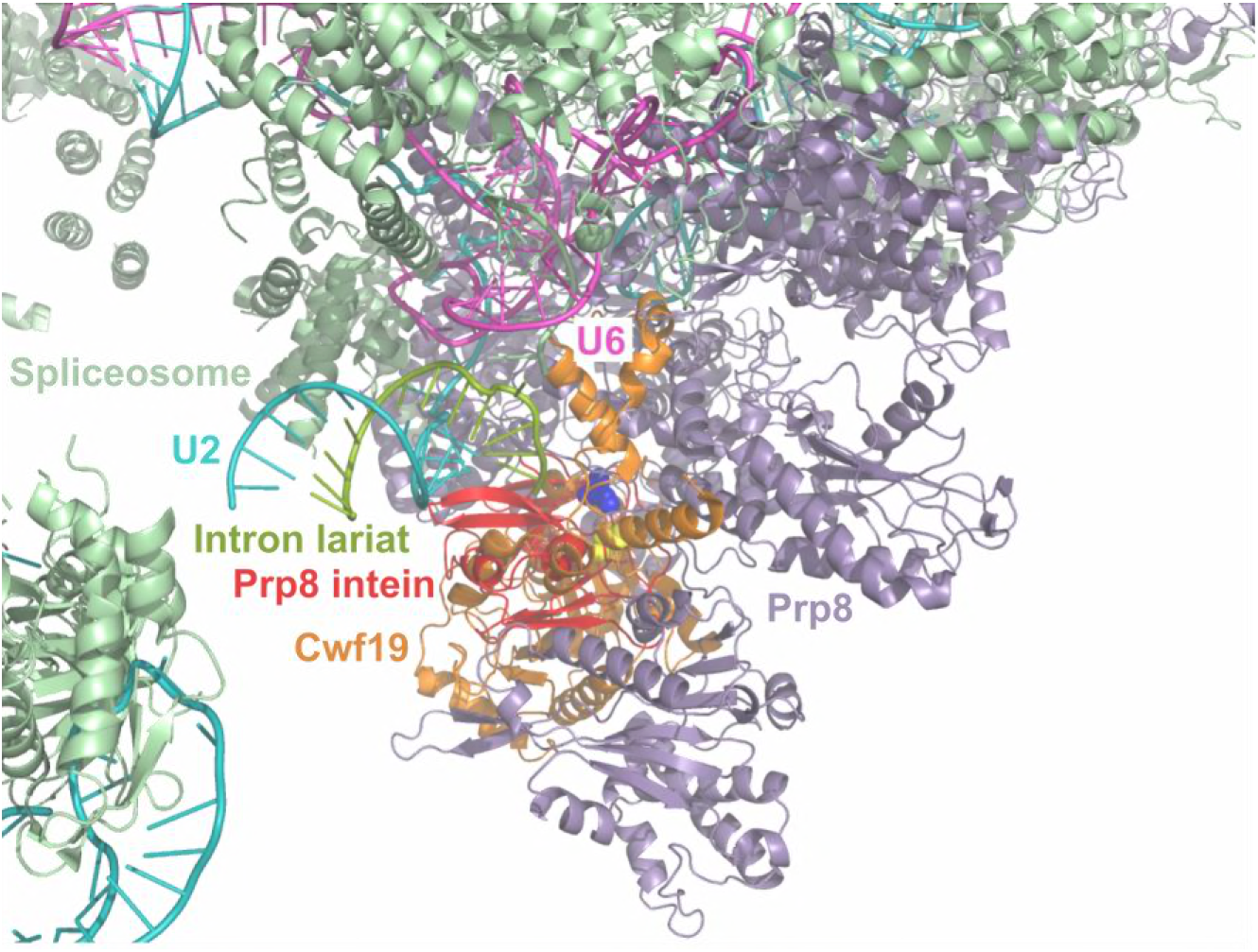
Model of the *Cne* Prp8 intein interrupting Prp8 and the spliceosome. The Prp8 intein-containing Prp8 precursor model was docked into a spliceosome structure from *Schizosaccharomyces pombe* (PDB 3JB9) to look for intein-spliceosome disruptions. Prp8 is shown as lavender and the Prp8 intein is shown as red, and the rest of the spliceosome is mint green. This reveals that the Prp8 intein would occupy a relatively sparse location in the spliceosome. The intein clashes are shown here (with labels) and noted in Fig. 6B.

**Table S1.** Bacterial strains

| Strain | Features and comments | Source |
| --- | --- | --- |
| DH5α | F<sup>-</sup> *endAI recAI hsdR17* (rK<sup>-</sup> mK<sup>-</sup>) *deoR supE44 thi-J gyrA96 relA* | Gibco-BRL |
| MG1655(DE3) | F<sup>-</sup> (λDE3) *ilvG rfb50 rph1* | James Imlay |
| BL21(DE3) | F<sup>-</sup> *ompT hsdS<sub>B</sub>*(r<sub>B</sub><sup>-</sup> m<sub>B</sub><sup>-</sup>) *gal dcm* (λDE3) | Novagen |

**Table S2.** Plasmids and constructs

| Plasmid/construct | Features and comments | Source |
| --- | --- | --- |
| pACYCDuet-1 | Expression vector, T7 promoter, Cam<sup>R</sup>. | Novagen |
| pACYC MIG SufB | Used as cloning backbone by removing *M. tuberculosis* SufB intein insert using ClaI/SphI. | Topilina et al. 2015 |
| pACYC MIG Prp8 | *C. neoformans* var. *grubii* H99 Prp8 intein flanked by short native exteins (N-extein: FWEKA; C-extein: SGFEE) cloned into ClaI/SphI sites between MBP and GFP coding sequences in pACYCDuet-1 backbone. | Present study |
| pACYC MIG Prp8 A-1V | Same as pACYC MIG Prp8 but with the indicated amino acid mutation at the last amino acid of the N-extein (position -1). | Present study |
| pACYC MIG Prp8 A-1V C61A/S/V | Same as pACYC MIG Prp8 A-1V but with the indicated amino acid mutation at the B block C61. | Present study |
| pACYC MIG *Afu* Prp8 | *Aspergillus fumigatus* 293 Prp8 intein flanked by short native exteins (N-extein: FWERA; C-extein: SGFEE) cloned into ClaI/SphI sites between MBP and GFP coding sequences in pACYCDuet-1 backbone. | Present study |
| pACYC MIG *Bde* Prp8 | *Batrachochytrium dendrobatidis* JEL423 Prp8 intein flanked by short native exteins (N-extein: FWEKA; C-extein: SGFEE) cloned into ClaI/SphI sites between MBP and GFP coding sequences in pACYCDuet-1 backbone. | Present study |
| pACYC MIG *Hca* Prp8 | *Histoplasma capsulatum* G186A Prp8 intein flanked by short native exteins (N-extein: FWERA; C-extein: SGFEE) cloned into ClaI/SphI sites between MBP and GFP coding sequences in pACYCDuet-1 backbone. | Present study |
| pET28a | Expression vector, T7 promoter, C-terminal His<sub>6</sub> tag, thrombin cleavage, Kan<sup>R</sup>. | Novagen |
| pET28a *Cne* Prp8 intein | *C. neoformans* var. *grubii* H99 Prp8 intein with 2 | Present |
|  | native N-extein residues (KA) in pET28a backbone at NcoI/XhoI sites. | study |
| pET47b | Expression vector, T7 promoter, N-terminal His<sub>6</sub> tag, PreScission cleavage, Kan<sup>R</sup>. | Novagen |
| pET47b *Cne* Prp8 intein 3 N-exteins | *C. neoformans* var. *grubii* H99 Prp8 intein with 3 native N-extein residues (EKA) in pET47b backbone at BamHI/NotI sites. | Present study |
| pET47b *Cne* Prp8 intein 3 N-exteins C1A/C61A/H65A | *C. neoformans* var. *grubii* H99 Prp8 intein with 3 native N-extein residues (EKA) plus C1A/C61A/H65A proposed binding site knockout in pET47b backbone at BamHI/NotI sites. | Present study |
| pET47b *Cne* Prp8 intein 3 N-exteins C1A | *C. neoformans* var. *grubii* H99 Prp8 intein with 3 native N-extein residues (EKA) and a single C1A mutation in pET47b backbone at BamHI/NotI sites. | Present study |
| pET47b *Cne* Prp8 intein 3 N-exteins C61A | *C. neoformans* var. *grubii* H99 Prp8 intein with 3 native N-extein residues (EKA) and a single C61A mutation in pET47b backbone at BamHI/NotI sites. | Present study |
| pET47b *Cne* Prp8 intein 3 N-exteins H65A | *C. neoformans* var. *grubii* H99 Prp8 intein with 3 native N-extein residues (EKA) and a single H65A mutation in pET47b backbone at BamHI/NotI sites. | Present study |

**Table S3.**
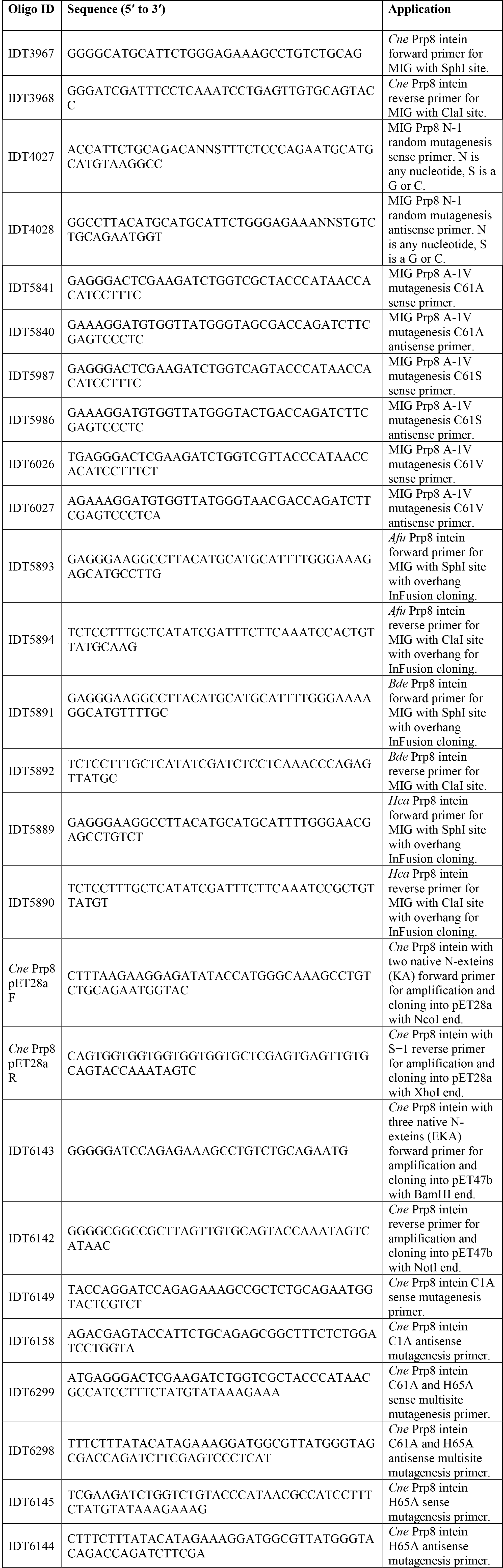
Oligonucleotide primers

**Table S4.** Data collection, refinement statistics, and model details for *Cne* Prp8 intein crystal structure

|  |  |
| --- | --- |
| ***Data Collection*** |  |
| Wavelength (Å) | 0.97946 |
| Resolution (Å) | 32.7-1.75 (1.81-1.75) |
| Space group | P 1 |
| Unit cell |  |
| a,b,c (Å) | 63.15 63.23 80.20 |
| $\alpha,\beta,\gamma$ (°) | 103.32 95.53 117.34 |
| Total reflections | 211,948 (19,137) |
| Unique reflections | 90,641 (8,259) |
| Multiplicity | 2.3 (2.3) |
| Completeness (%) | 85.91 (77.71) |
| Mean $I/\sigma(I)$ | 12.48 (1.55) |
| $R_{merge}$ (%) | 0.069 (0.91) |
| ***Refinement*** |  |
| $R_{work}$ | 0.197 (0.293) |
| $R_{free}$ | 0.235 (0.310) |
| No. of non-H atoms | 7576 |
| macromolecules | 7026 |
| solvent | 550 |
| RMS (bonds) (Å) | 0.005 |
| RMS (angles) (°) | 1.02 |
| Ramachandran |  |
| favored (%) | 98.42 |
| allowed (%) | 1.58 |
| outliers (%) | 0.00 |
| Rotamer outliers (%) | 0.00 |
| Clashscore | 3.45 |
| Average B-factor (Å<sup>2</sup>) | 30.39 |
| macromolecules | 29.81 |
| solvent | 37.80 |

Statistics for the highest-resolution shell are shown in parentheses.

